# Single-cell RNA sequencing reveals mRNA splice isoform switching during kidney development

**DOI:** 10.1101/688564

**Authors:** Yishay Wineberg, Tali Hana Bar-Lev, Anna Futorian, Nissim Ben-Haim, Leah Armon, Debby Ickowicz, Sarit Oriel, Efrat Bucris, Yishai Yehuda, Naomi Pode-Shakked, Shlomit Gilad, Sima Benjamin, Peter Hohenstein, Benjamin Dekel, Achia Urbach, Tomer Kalisky

## Abstract

During mammalian kidney development, nephron progenitors undergo a mesenchymal to epithelial transition and eventually differentiate into the various tubular segments of the nephron. Recently, the different cell types in the developing kidney were characterized using the Dropseq single cell RNA sequencing technology for measuring gene expression from thousands of individual cells. However, many genes can also be alternatively spliced and this creates an additional layer of heterogeneity. We therefore used full transcript length single-cell RNA sequencing to obtain the transcriptomes of 544 individual cells from mouse embryonic kidneys. We first used gene expression levels to identify each cell type. Then, we comprehensively characterized the splice isoform switching that occurs during the transition between mesenchymal and epithelial cellular states and identified several putative splicing regulators, including the genes Esrp1/2 and Rbfox1/2. We anticipate that these results will improve our understanding of the molecular mechanisms involved in kidney development.

## INTRODUCTION

Kidney development is a complex process that involves multiple interacting cell types [1–5]. It starts at the embryonic stage and continues from week 5 to week 36 of gestation in humans and from day E10.5 to approximately day 3 after birth in mice. The process is initiated by signaling interactions between two lineages originating from the intermediate mesoderm - the ureteric duct and the metanephric mesenchyme (Fig. S1). These interactions invoke the ureteric duct to invade the metanephric mesenchyme creating a tree-like structure. Around the tip of each branch of this tree, the “ureteric tip”, cells from the metanephric mesenchyme are induced to condense and form the “cap mesenchyme”, which is a transient nephron progenitor cell population (NPCs). Next, cells from the cap mesenchyme undergo a mesenchymal to epithelial transition (MET) and progressively differentiate into early epithelial structures: pretubular aggregates, renal vesicles, and comma and S-shaped bodies. The S-shaped bodies further elongate and differentiate to form the various epithelial tubular segments of the fully developed nephron, whose main constituents are the podocytes, the proximal tubule, the loop of Henle, and the distal tubule. At an early stage in their differentiation the distal tubules connect to the ureteric tips that form the collecting duct system for draining the nephrons. Meanwhile, the un-induced cells of the metanephric mesenchyme differentiate into other supporting cell types of the kidney such as interstitial fibroblasts, pericytes, and mesangial cells (Fig. S1). In the last few years the various cell populations of the developing kidney were characterized [6–8], mainly using the Dropseq single cell RNAseq protocol that enables measuring of gene expression levels from many thousands of individual cells [9].

A central process in kidney development is the mesenchymal to epithelial transition (MET) that occurs during the differentiation of cells from the metanephric mesenchyme to the cap mesenchyme and then to nephron tubules. Similar transitions from mesenchymal to epithelial states and vice-versa (Epithelial-Mesenchymal Transition, EMT) are thought to play a central role in development, as well as in pathological processes such as cancer metastasis [10] and organ fibrosis [11]. There are significant structural and functional differences between mesenchymal and epithelial cells: while mesenchymal cells are typically loosely associated with each other, surrounded by an extracellular matrix, and have migratory capabilities, epithelial cells are tightly interconnected by junctions and are polarized with distinct apical and basolateral membranes. Thus, epithelial cells can create 2-dimensional surfaces and tubes with a clear in/out distinction that are capable of absorption and secretion. There are also large differences in gene expression: mesenchymal cells typically express Fibronectin (Fn1), Vimentin (Vim), and N-cadherin (Cdh2) [12], as well as the transcription factors Snai1/2, Zeb1/2, and Twist1/2, while epithelial cells typically express other genes such as E-cadherin (Cdh1) and Epcam.

It was recently realized that mesenchymal and epithelial cells also express alternative splice isoforms of genes that are expressed in both cell states. For example, in many systems, both in-vivo and in-vitro, the genes Enah, Cd44, Ctnnd1, and Fgfr2 were found to be expressed in both mesenchymal and epithelial states, but with unique isoforms specific to each state [13–18]. It was also found that RNA binding proteins such as ESRP1/2, RBFOX1/2, RBM47, QKI, and others act as splicing regulators that promote splicing of specific mesenchymal or epithelial variants [12,17,19,20]. mRNA splicing creates an additional layer of heterogeneity that, apart from specific genes [21–23], has not yet been comprehensively studied in the developing kidney.

Therefore, in this study we set to characterize the splice isoform switching events that occur during the transition between the mesenchymal and epithelial cellular states in the course of kidney development by comparing gene expression and alternative splicing in the various mesenchymal and epithelial cell populations. Since the kidney is a heterogeneous organ that is composed of numerous cell populations of widely varying proportions [6–8,24], it is difficult to isolate pure populations of mesenchymal and epithelial states. Moreover, typical sequencing depths from a single cell are not sufficient for splicing analysis. We therefore performed single cell RNA sequencing on 576 individual cells from the kidneys of E18.5 mouse embryos using the Smartseq2 protocol for sequencing full transcript lengths [25,26]. We first identified the main cell lineages that coexist in the nephrogenic zone of the fetal developing kidney – the uninduced metanephric mesenchyme, the cap mesenchyme, podocytes, epithelial cells, endothelial cells, and infiltrating immune cells (e.g. macrophages). We then merged the raw reads from all cells belonging to each population in order to create “bulk” *in-silico* transcriptomes that represent each cell population. These “bulk” transcriptomes had sufficient sequencing depth to allow us to characterize splice isoform switching and to identify putative splicing regulators.

## RESULTS

### Gene expression levels were used to classify each cell into one of the various cell types that coexist within the nephrogenic zone of the developing mouse fetal kidney

We collected kidneys from transgenic mouse embryos that express GFP under the control of a Six promoter [27] (Fig. 1A). These mice have the advantage that cells from the cap mesenchyme – previously shown to express the transcription factor Six2 - are fluorescently tagged and can be enriched by flow cytometry. The kidneys were collected at day E18.5 of gestations since at this stage we expect to observe a still-active nephrogenic zone containing both early progenitor populations as well as fully developed nephrons [6,7]. After kidney dissociation, we used flow cytometry to select 288 cells expressing high levels and 288 cells expressing low levels of the Six2-GFP transgene, and for each individual cell we performed full transcript length single-cell RNA sequencing using the Smartseq2 protocol [25,26]. After discarding low quality cells, this resulted in gene expression and sequence information for 544 individual cells, with approximately equal proportions of cells originating from the Six2-high and Six2-low fractions.

**Figure 1:**
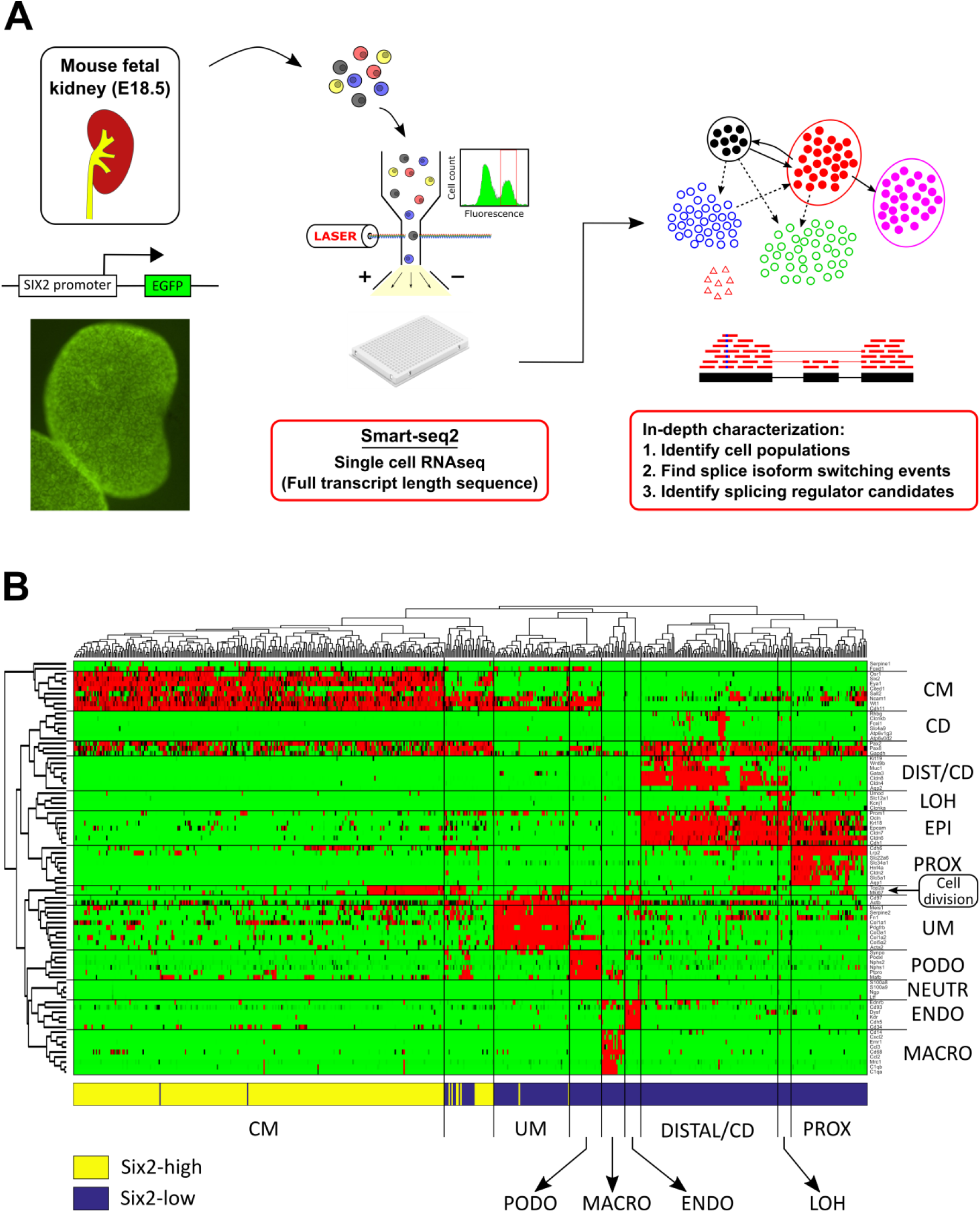
We used single cell RNA sequencing to identify and transcriptionally characterize the main cell lineages that coexist in the nephrogenic zone of the developing mouse fetal kidney. (A) Shown is the general outline of the experiment. Kidneys from transgenic mouse embryos with a Six2-GFP reporter gene were harvested at day E18.5. The Six2-GFP reporter gene shows a clear fluorescent pattern marking the cap mesenchyme. After tissue dissociation, single cells from the Six2-high and Six2-low cell fractions were sorted into individual wells and their mRNA was sequenced for full transcript length using the Smartseq2 protocol. (B) A clustergram for 83 selected genes from the literature vs. 544 cells shows the main cell lineages in the developing kidney. We manually classified the cells to lineages known to co-exist in the nephrogenic zone of the developing fetal kidney, including the un-induced mesenchyme (UM), the cap mesenchyme (CM), podocytes (PODO), proximal tubular epithelial cells (PROX), the loop of Henle (LOH), distal tubular cells and collecting duct (DIST/CD), endothelial cells (ENDO), and infiltrating immune cells - mainly macrophages (MACRO) and a small number of neutrophils (NEUTR). The horizontal bar at bottom of the figure depicts the gate used by FACS to sort each cell (Six2-high or Six2-low). Consistent with the fact that Six2 is highly expressed in the cap mesenchyme (CM), it can be seen that cells originating from the Six2-high fraction predominantly belong to the cap mesenchyme (CM), whereas cells originating from the Six2-low fraction belong mostly to the other populations. The gene panel includes genes that were shown to be specific to the different populations, as well as general epithelial markers (EPI) and genes that are over-expressed in the S-G2-M phase of the cell cycle, indicating cell division. Notice that some cells (within the 2^nd^ cluster from the left) express a mixture of cap mesenchyme (CM) markers and epithelial markers (EPI, PODO), as well as cell division. These are presumably cells from early epithelial structures - presumably pre-tubular aggregates, renal vesicles, and C/S-shaped bodies; see Supplementary Information.

Using expression levels of selected genes from the literature that were shown to be specific to each population (e.g. [6,7]), as well as general epithelial markers and genes indicating cell division, we manually classified each cell into one of the major cell types that co-exist in the nephrogenic zone of the developing kidney (Figs. 1B, 2A, S1, S4-S9). These include the un-induced mesenchyme (UM), the cap mesenchyme (CM), podocytes (PODO), early epithelial structures (PROX_1) - presumably pre-tubular aggregates, renal vesicles, and C/S-shaped bodies; proximal epithelial tubules (PROX_2), the loop of Henle (LOH), distal tubule and collecting duct (DIST/CD), endothelial cells (ENDO), and infiltrating immune cells, mainly macrophages (MACRO). PROX_1 cells differ from PROX_2 cells in that they over-express markers for early epithelial structures such as Mdk and Lhx1 (Fig. S7) as well as markers for actively dividing cells such as Mki67 and Top2a (Fig. S10). We note that we found it extremely difficult to distinguish between cells of the distal tubule and cells of the collecting duct in our dataset, probably due to their transcriptional similarity as well as the relatively small number of cells in this experiment, and therefore we merged them into a single population.

**Figure 2:**
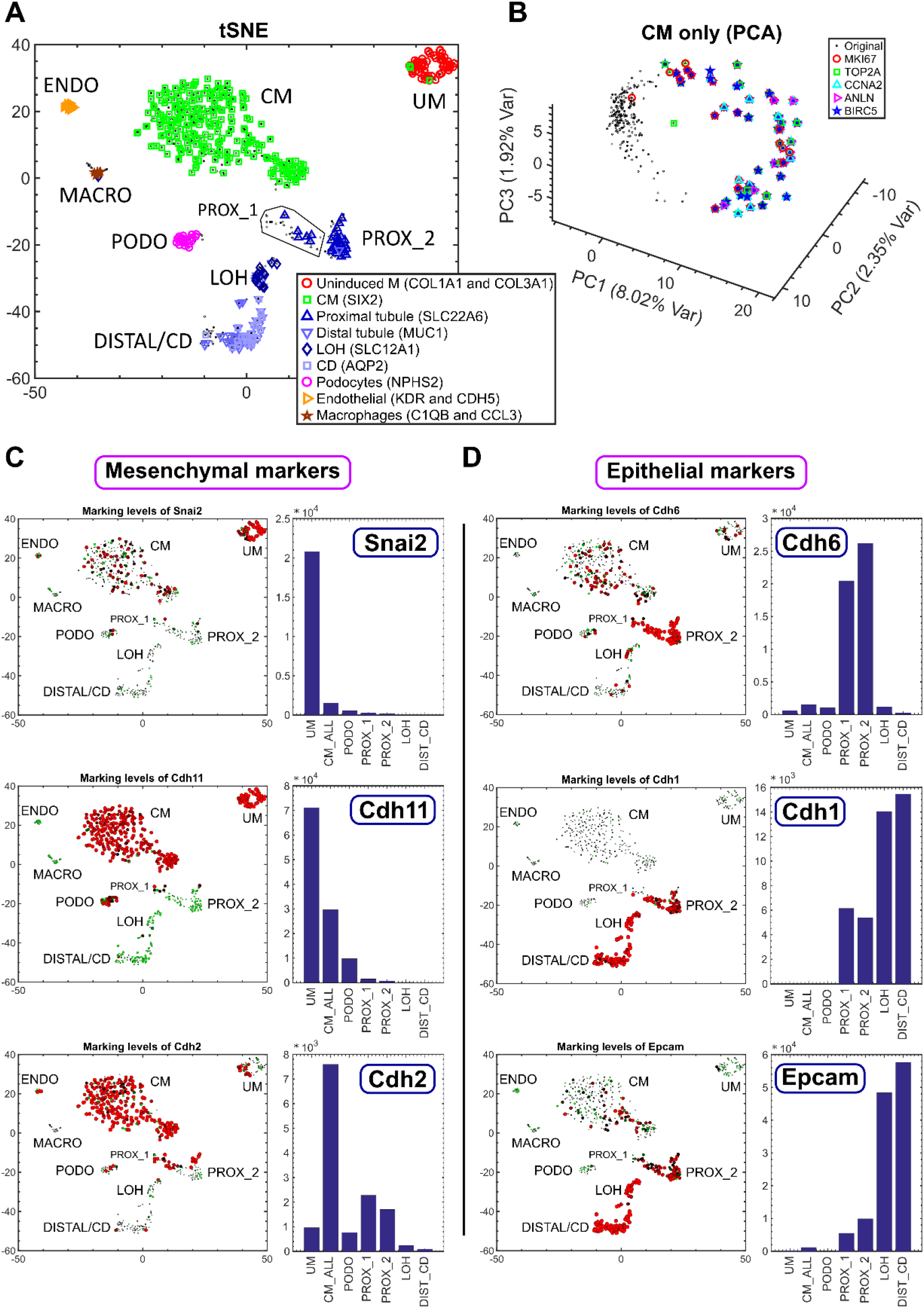
Single-cell gene expression analysis enables characterization of cellular heterogeneity, cell cycle dynamics, and the Mesenchymal to Epithelial Transition (MET) in the developing mouse fetal kidney. (A) Shown is a tSNE plot of 544 single cell gene expression profiles, each consisting of 728 highly variable genes (see Methods). Each cell is represented by a dot. Cells over-expressing genes that were previously shown to mark different cell types are marked by additional symbols. UM – un-induced mesenchyme, CM – Cap mesenchyme, PODO – podocytes, PROX_1 – early epithelial structures (presumably pre-tubular aggregates, renal vesicles, and C/S-shaped bodies), PROX_2 – proximal epithelial tubules, DIST/CD – distal tubules and collecting duct, ENDO – endothelial cells, MACRO – macrophages. (B) Cells of the Cap mesenchyme create a circular manifold in gene expression space that corresponds to the cell cycle. Shown is a PCA figure of cells from the CM only. Each cell is represented by a dot. Cells over-expressing genes such as Top2a and Mki67 – genes that were shown to be over-expressed in the S-G2-M phases of the cell cycle - are marked by additional symbols. These cells are located in a specific segment of the circular manifold representing the S-G2-M segment of the cell cycle. (C, D) Expression levels of the mesenchyme related genes Snai2, Cdh11, and Cdh2 are typically higher in the UM and CM, whereas and the epithelial genes Cdh6, Cdh1, and Epcam are typically higher in the PROX_1, PROX_2, LOH, and DIST_CD. Shown are tSNE plots and barplots showing the expression levels of selected genes. The area of each circle in each tSNE plot is proportional to log2(1+expression) of the specific gene in that particular cell. The expression level in each cell is also encoded by the circle color (red – high expression, green - low expression). The barplots show gene expression levels in “bulk” *in-silico* cell transcriptomes representing the different populations (UM, CM, PODO, PROX_1, PROX_2, LOH, and DIST_CD) that were created by uniting raw reads from all cells belonging to each population. The annotations “CM_ALL” and “CM” are used interchangeably to represent all cells that were classified as belonging to the cap mesenchyme (see also Supplementary Information).

After identifying the various cell subpopulations, we inspected the expression levels of the genes Mki67 and Top2a that are known to be over-expressed during cell division (Fig. S10). We found that the un-induced mesenchyme (UM), the cap mesenchyme (CM), the early epithelial structures (PROX_1), the loop of Henle (LOH), and distal tubular cells/collecting duct (DIST_CD) each contain a substantial subset of dividing cells that over-express these genes, whereas the podocytes (PODO) and proximal epithelial tubules (PROX_2) do not. Moreover, we found that the cells of the cap mesenchyme create a circular manifold (i.e., a high dimensional ring) in gene expression space, whose segments correspond to the different phases of the cell cycle (Figs. 2B, S11). Using RNA velocity [28] - a computational tool for inferring a vector between the present and predicted future transcriptional state of each single cell by distinguishing between the spliced mRNA (present state) and yet-unspliced mRNA (future state) - we observed a consistent directional flow along this circular manifold (Fig. S11).

Next, we inspected the expression levels of genes that are known to be involved in the Mesenchymal to Epithelial Transition (MET) or that are known to be preferentially over-expressed in mesenchymal or epithelial lineages [10,12,29] (Figs. 2C-D, S12, S13). We observed high levels of mesenchyme-associated genes such as Snai2, Cdh11, and Cdh2 in the earlier developmental lineages – the un-induced mesenchyme (UM) and cap mesenchyme (CM). Likewise, higher levels of epithelial genes such as Cdh6, Cdh1, and EpCAM were prevalent in the more differentiated lineages – the early epithelial structures (PROX_1), proximal tubules (PROX_2), loop of Henle (LOH), and the distal tubular cells and collecting duct (DIST/CD). We noticed that the expression of Cdh11 showed a gradual decrease (Figs. 2C, S12), with the highest expression levels being expressed in the un-induced mesenchyme (UM), medium levels in the cap mesenchyme (CM), even lower levels in the podocytes (PODO), and very low levels in the epithelial lineages (PROX_1 and PROX_2). Likewise, when comparing Cdh6 and Cdh1 (Figs. 2D, S12) we noticed that Cdh6 is higher in the early epithelial structures (PROX_1) and proximal tubules (PROX_2) whereas Cdh1 is higher in the loop of Henle (LOH) and the distal tubules/collecting duct (DIST/CD) [29].

### rMATS was used to characterize the splice isoform switching events that occur during the transition between the mesenchymal and epithelial cellular states in the course of kidney development

Since we obtained full transcript length sequence information, we were able to use rMATS [30] to characterize the splice isoform switching that occurs during the mesenchymal to epithelial transition (MET) in the course of kidney development. In particular, we focused on identifying alternatively spliced exons (cassette exons) by searching for exons whose inclusion levels – defined as the fraction of transcripts that include the exon out of the total number of transcripts that either include the exon or skip over it - changes significantly between the mesenchymal and epithelial states. Since the coverage for each single cell was rather low for splicing analysis, we first merged the raw reads from all cells belonging to each population in order to create “bulk” *in-silico* transcriptomes that represent each cell population, and then used the resulting “bulk” transcriptomes as input to rMATS. We searched for cassette exons whose inclusion levels change significantly (FDR=0, difference in inclusion levels > 0.2) between either of the mesenchymal populations – the un-induced mesenchyme (UM) or the cap mesenchyme (CM) and all of the epithelial populations - the early epithelial structures (PROX_1), proximal tubule (PROX_2), loop of Henle (LOH), and distal tubule/collecting duct (DIST/CD).

We found a list of 57 cassette exons that were thus differentially expressed between the mesenchymal and epithelial lineages (Figs. 3A, 3C). These exons include some known examples that were previously observed in EMT such as the epithelial-associated cassette exons in Map3k7 [31–33], Dnm2 [31,34,35], Pard3 [34], and the mesenchymal-associated exons Plod2 [12,18,36], Csnk1g3 [18,33], and Ctnnd1 [14,18]. Gene Ontology (GO) enrichment analysis (Fig. 3B, Table S3) showed that the genes containing these exons are related to epithelial characteristics (e.g. cell-cell junction and Cdh1 interactions) or mesenchymal characteristics (e.g. lamellipodium or cell leading edge – related to cellular motility).

**Figure 3:**
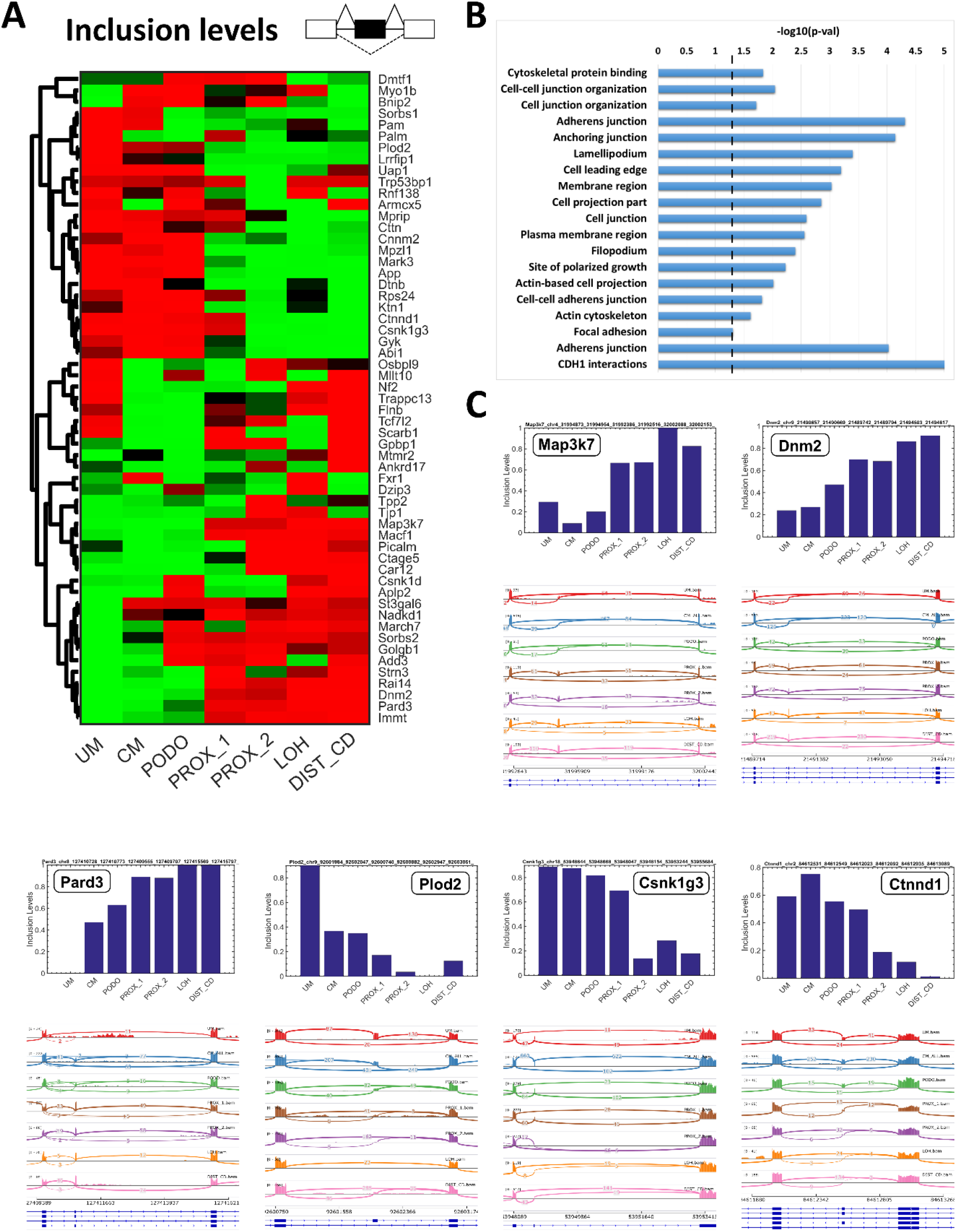
Characterization of splice isoform switching events that occur during the Mesenchymal to Epithelial transition (MET) in the course of kidney development. (A) Shown is a heatmap of inclusion levels of selected cassette exons that change significantly (FDR=0, Difference in inclusion levels > 0.2) between either of the mesenchymal populations (UM or CM) and all the epithelial populations (PROX_1, PROX2, LOH, and DIST_CD). Exon inclusion levels were derived from *in-silico* “bulk” transcriptomes representing the different cell populations (UM, CM, PODO, PROX_1, PROX_2, LOH, and DIST_CD) that were created by uniting raw reads from all cells belonging to each population. Colors indicate relative high (=red) vs. low (=green) inclusion levels. The inclusion levels for each exon (=row) were independently standardized by mean-centering and dividing by the standard deviation. (B) Gene Ontology (GO) enrichment analysis shows that the genes containing these differentially expressed cassette exons are related to structural and functional properties of epithelial cells (e.g. cell-cell junctions and CDH1 interactions) or to characteristics of mesenchymal cells (mainly cellular motility, e.g. lamellipodium and cell leading edge). (C) Sashimi plots and Barplots of inclusion levels in selected exons show gradual increase (Map3k7 [31–33], Dnm2 [31,34,35], and Pard3 [34]) or decrease (Plod2 [12,18,36], Csnk1g3 [18,33], and Ctnnd1 [14,18]) of inclusion levels during the transition from mesenchymal (UM and CM) to epithelial states (PROX_1, PROX_2, LOH, and DIST_CD).

### Supervised analysis identifies additional genes that undergo splice isoform switching during kidney development

Since the low coverage and bias of single-cell protocols limits the power of automated tools for discovering splice isoform switching, we searched the existing literature for additional genes for which different splice isoforms are expressed in mesenchymal versus epithelial cells or which are known to undergo splice isoform switching during EMT, and examined their alternative splicing between the different cell populations of the developing kidney. Indeed, we found that the genes Fgfr2 [14,37] (Fig. S15), Epb41l5 [17,35] (Fig. S15), Fat1 [18,34] (Fig. S16), and Arhgef10l [19,20,38] (Fig. S17) express their mesenchymal isoforms predominantly in the early mesenchymal populations (UM and CM), and their epithelial isoforms mostly in the more differentiated epithelial populations (LOH and DIST/CD). The early epithelial structures (PROX_1) and the proximal tubules (PROX_2) express either the mesenchymal or the epithelial isoform or a mixture of both. Interestingly, we observed that the podocytes (PODO) in many cases express the mesenchymal rather than the epithelial isoforms (Figs. 3A, S15-17). This reveals another aspect in which podocytes, which form a specialized type of epithelial tissue [5], are different from most other forms of epithelia.

The genes Enah and Cd44 are prominent examples of genes that undergo splice isoform switching during EMT [17,18]. Both genes contain cassette exons that are expressed in epithelial cells only. However, this behavior was impossible to observe in our single-cell dataset, probably due to the relative low coverage and bias of our single-cell protocol with respect to “bulk” RNA sequencing. We therefore performed additional “bulk” RNA sequencing on three replicates of sorted Six2-high and Six2-low cell fractions. Since Six2 is uniquely expressed in the cap mesenchyme (CM), the cell fraction that was gated Six2-high is predominantly composed of cells originating from the cap-mesenchyme, whereas the cell fraction gated for Six2-low contains a mixture of all the other mesenchymal and epithelial populations. Nevertheless, by manually comparing Sashimi plots from these two cell fractions we were able to observe the alternatively spliced cassette exons in the genes Enah and Cd44 (Fig. S18).

A different form of alternative splicing, not related to EMT, was observed in gene Cldn10, which is an important component of epithelial tight junctions in the kidney and provides a barrier and permits selective para-cellular transport [39]. Two isoforms of the gene Cldn10 were previously found to co-exist in the kidney, one being highly expressed in the cortex and the other in the medulla [40]. These alternatively spliced isoforms are thought to generate different permselectivities in different segments of the nephron. In our dataset we observed that the cortical isoform was indeed over-expressed in the early epithelial structures (PROX_1) and the proximal tubules (PROX_2) - which are located in the cortex - while the medullary isoform was predominantly expressed in the loop of Henle (LOH) which is predominantly located in the medulla (Fig. S19). This confirmed the previous in-situ hybridization measurements [40] also at the single-cell transcriptomic level.

We also inspected the Wt1 gene, which is essential for normal kidney development [41–44]. Mutations and alternative splicing in Wt1 were found to play an important role in developmental defects such as Denys-Drash syndrome and Frasier syndrome, as well as in Wilms’ tumors. Wt1 is known to encode for multiple possible splice isoforms [41,45]: for example, three amino acids (K-T-S) at the 3’ end of exon 9 may be included, creating a KTS+ isoform, or skipped, resulting in a KTS− isoform (Fig. S20). It was previously found that these isoforms differ in their affinity to DNA [46] and in their localization to different compartments within the nucleus [47]. Moreover, it was found that normal tissues have a KTS+:KTS− ratio of approximately 0.6, while in tissue from patients with Frasier syndrome the amount of KTS+ transcripts decreases, resulting in a lower KTS+:KTS− ratio of approximately 0.4 [42,48,49]. In our dataset we observed a decrease in the KTS+:KTS− isoform ratio, starting from from ~0.75 in the un-induced mesenchyme (UM) and the cap mesenchyme (CM), and converging to ~0.6 in the podocytes (PODO), early epithelial structures (PROX_1), and proximal tubules (PROX_2) (Supplementary Information and Fig. S20). It was interesting to observe the relative similarity of the KTS+:KTS− ratio in the podocytes and epithelial populations (PODO, PROX_1, and PROX_2), which was lower than the ratio for the mesenchymal populations (UM, CM). This might indicate the existence of a mechanism for stabilizing and tightly regulating this ratio in maturing renal cell populations. Likewise, we also observed a gradual increase in the inclusion levels of cassette exon 5 (Fig. S20).

We note that when inspecting the expression behavior of Wt1 we found multi-level expression differences between the different cell populations (Fig. S20). Wt1 is most highly expressed in the podocytes [50] (PODO), in which it was previously shown to be a key transcriptional regulator [44], moderately expressed in the un-induced mesenchyme (UM, partially) and cap-mesenchyme (CM), and under-expressed in the loop of Henle (LOH) and distal tubules/collecting duct (DIST/CD). In the early epithelial structures (PROX_1) we observed a wide distribution of Wt1 expression, probably due to the fact that some cells (e.g. those in the cleft of the S-shaped body [5]) are in the process of differentiating to podocytes while others are destined to become constituents of the proximal tubule, loop of Henle, or distal tubule.

### Differential expression of RNA binding proteins and RNA binding motif enrichment analysis suggest that Esrp1/2 and Rbfox1/2 are splicing regulators of the Mesenchymal to Epithelial Transition (MET) that occurs during kidney development

We next used rMAPS [29] to identify putative RNA binding proteins (RBP’s) that act as splicing regulators for splice isoform switching between the mesenchymal and epithelial states during renal development. We first compared the mean expression levels of 84 known RNA binding proteins [51–53] between the mesenchymal (UM, CM) and epithelial populations (PROX_1, PROX_2, LOH, DIST_CD) and found several putative splicing regulators that were differentially expressed (Figs. 4A, S21).

**Figure 4:**
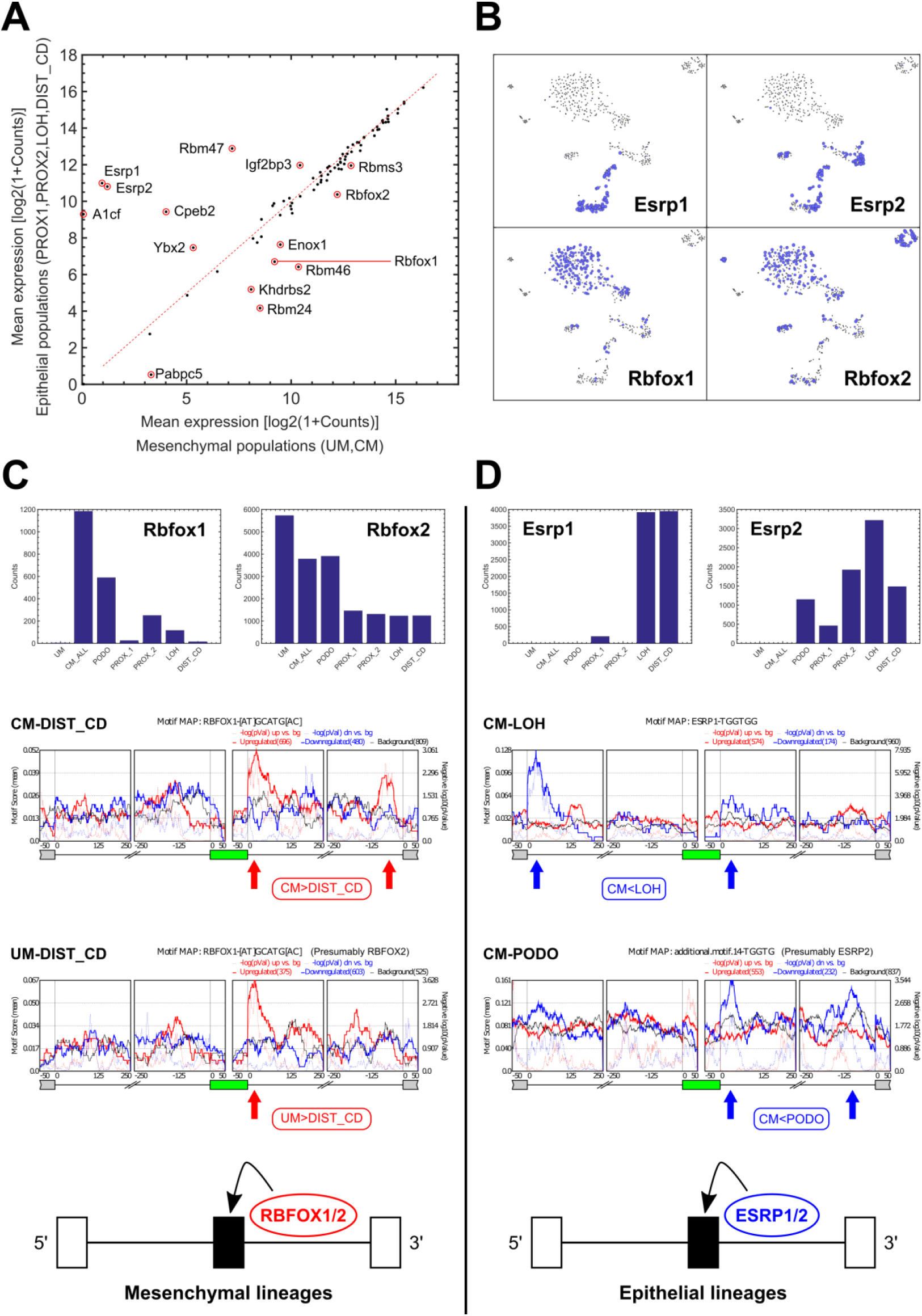
Differential expression of RNA binding proteins and RNA binding motif enrichment analysis suggest that Esrp1/2 and Rbfox1/2 are splicing regulators of the Mesenchymal to Epithelial Transition (MET) that occurs during kidney development. (A) Shown is a comparison between mesenchymal and epithelial states of the mean expression levels of 84 RNA binding proteins (RBP’s) that are known from the literature to regulate splicing through binding of mRNA transcripts. We used expression levels from “bulk” *in-silico* transcriptomes that represent each cell population. (B) A tSNE plot of the single cell profiles, highlighting cells that express the putative splicing regulators RBFOX1/2 and ESRP1/2. It can be seen that RBFOX1 and RBFOX2 are highly expressed in the mesenchymal populations, while ESRP1 and ESRP2 are highly expressed in the epithelial populations. The area of each circle is proportional to log2(1+expression) of each gene in that particular cell. (C,D) Cassette exons that are over-expressed in the mesenchymal populations contain a significant enrichment of RBFOX1/2 binding motifs at their downstream introns [12,52,54]. Likewise, cassette exons that are over-expressed in the epithelial populations contain a significant enrichment of ESRP1/2 binding motifs in their downstream introns [19,31,38], and in some cases (CM vs. LOH), also in the far 5’ end of their upstream introns (as previously observed in EMT [31]). This indicates that RBFOX1/2 and ESRP1/2 are splicing regulators [12] involved in the Mesenchymal to Epithelial Transition (MET) that occurs during kidney development. The annotations “CM_ALL” and “CM” are used interchangeably to represent all cells that were classified as belonging to the cap mesenchyme; see also Methods and Supplementary Information.

Of these differentially expressed RNA binding proteins, we found Rbfox1 and Rbfox2 to be over-expressed in the mesenchymal cells, while Esrp1 and Esrp2 are over-expressed in the epithelial cells (Fig. 4B). Likewise, we found that Rbfox1/2 and Esrp1/2 have RNA binding sites (motifs) that are enriched in the 5’ or 3’ neighboring introns of the cassette exons that are differentially expressed between mesenchymal and epithelial states (Figs. 4C-D, S22) [12,19,31,38]. This indicates that RBFOX1/2 and ESRP1/2 are splicing regulators involved in Mesenchymal to Epithelial Transition (MET) during kidney development, similar to what was previously observed in EMT [12]: in the mesenchymal cell populations Rbfox1 and/or Rbfox2 bind to mRNA in the downstream 3’-flanking introns of the mesenchymal associated cassette exons and promote their inclusion [12,52,54], while in the epithelial cell populations Esrp1 and/or Esrp2 bind to the downstream 3’-flanking introns [19,31,38] (and in some cases also to an additional site at the far 5’ end of the upstream flanking intron [31]) of the epithelial associated cassette exons and promote their inclusion.

## DISCUSSION

In this study we used the Smartseq2 protocol [25,26] for full transcript length single-cell RNA sequencing to characterize the splice isoform switching events that occur during the Mesenchymal to Epithelial Transition (MET) in the course of kidney development. Such splicing information is not obtainable using 3’-end digital counting protocols such as Dropseq [6–9], apart from splicing events located at the very 3’ end of mRNA transcripts. These results highlight the importance of combining 3’-end digital counting technologies for transcriptional profiling of many thousands of cells, with full transcript length RNA sequencing for deeper analysis of selected cell populations, in order to obtain a detailed understanding of the molecular mechanisms involved in kidney development and disease.

Since the Smartseq2 protocol that we used is practically limited to a few hundreds of cells, we were unable to detect very small immune populations [55] or to discern between all cell subtypes. For example, we were unable to discern subpopulations within the very early epithelial structures (PROX_1) or stromal subtypes within the uninduced mesenchyme (UM) [6]. Nevertheless, the Smartseq2 protocol does have the advantage of being able to measure expression levels more precisely. For example, we were able to discern high, medium, and low expression levels of genes such as Cdh11 (Fig. S12) or Wt1 (Fig. S20).

Each cell in our analysis was sequenced at roughly 1-2 million reads per cell. Since for splicing analysis roughly 20-40 million reads are typically required, we merged the raw reads from all cells belonging to each population in order to create “bulk” *in-silico* transcriptomes that represent each cell population and then performed splicing analysis on these. We note, however, that splicing analysis can also be done for individual cells [56], but due to the small number of reads the inferred inclusion levels for most genes will be less accurate and with wide margins of error except for the most highly expressed genes.

We note that the motif analysis for RNA binding proteins (RBP’s) often resulted in non-specific results, such as candidate splicing regulators that were not even expressed in some populations. We hypothesize that this stems from the fact that the binding motifs are not very specific since they are typically only a few bases long. We therefore based our identification of the splicing regulators Esrp1/2 and Rbfox1/2 on the existence of two additional criteria apart from RNA binding motif enrichment: first, the expression levels of Esrp1/2 and Rbfox1/2 differ significantly between the mesenchymal and epithelial cell states, and second, there is much previous evidence for similar functionality in other developing organs and in-vitro systems [12,14,17,31,32,35,38]. The marked differences in expression between mesenchymal and epithelial populations of other RNA binding proteins such as Cpeb2, Rbm47 [12], Msi1, Rbms3, and others (Figs. 4A, S21) indicate that they might also be involved in renal MET splicing regulation. However, we did not observe a consistent enrichment of known binding motifs for these genes as we did for Esrp1/2 and Rbfox1/2. Likewise, there may be additional splicing regulators whose expression does not change and whose RNA binding activity is modulated by protein modification. One example is QK (also known as QKI) [12], but we did not find significant motif enrichment for this gene in the MET associated differentially expressed cassette exons.

In a recent study it was shown that ablation of Esrp1 in mice, alone or together with Esrp2, resulted in reduced kidney, size fewer ureteric tips, reduced nephron numbers, and a global reduction of epithelial splice isoforms in the transcriptome of ureteric epithelial cells [57]. We believe that our results provide a detailed picture at the single-cell level that complements the above study. Moreover, the fact that kidneys still develop under the ablation of Esrp1/2, taken with our results, suggests that there are multiple splicing regulators acting combinatorically thus creating a bypass mechanisms so that one splicing regulator compensates, although partially, for lack of another. Since with current technology it is infeasible to create transgenic mice containing knockouts of the many possible combinations of multiple regulators, we suggest using kidney organoids as a model system along with single cell analysis for future functional studies.

## METHODS

### Tissue collection, dissociation and flow cytometry

A wild-type female mouse was crossed with a male that was heterozygous for the Six2-GFP transgene [27]. The female was sacrificed at day E18.5 of pregnancy and kidneys from eleven embryos were dissected, placed in PBS on ice, and examined under a fluorescent microscope to check for the presence of GFP. Kidneys from transgenic embryos showed a clear fluorescent pattern marking the cap mesenchyme (Fig. 1A), whereas those from the non-transgenic embryos showed uniform background fluorescence. Seven out of the eleven embryos contained the Six2-GFP transgene. Keeping the transgenic and non-transgenic kidneys separate, each kidney was then cut into 2-4 small pieces using a surgical razor blade and placed in 1ml of trypsin 0.25% (03-046-1A, Trypsin Solution B (0.25%), Biological Industries) using forceps. Initial tissue trituration was performed using a tissue grinder (D8938, Dounce, Large clearance pestle) followed by up-down pipetting with a P1000 pipette. Tissue fragments were then incubated with trypsin at 37°C for 20 minutes, followed by additional trituration by pipetting. After visual confirmation that the majority of cells were fully dissociated, enzyme digestion was stopped by adding 2ml of DMEM with 10% FBS and placed on ice. Cells were first filtered with 70 micron cell strainers (CSS-010-070, Lumitron) and then 40 micron cell strainers (732-2757, VWR), pelleted by centrifugation for 5 minutes at 500 x g, re-suspended in 3ml PBS, and kept on ice until FACS sorting.

Single cell sorting was done using a BD FACSAria III flow cytometer with an 85 micron nozzle. Forward and side scatter were used to filter out red blood cells and to select for live single cells. Gating based on the negative control – the cells from the non-transgenic kidneys - was used to select for cells that were either positive (Six2-high) or negative (Six2-low) for expression of the Six2-GFP transgene.

All procedures were approved by the institutional Animal Care and Use Committee in Bar-Ilan University.

### Single cell RNA sequencing

For single cell RNA sequencing, single cells were sorted into 96 individual wells from a 384 well plate that were pre-filled with Smartseq2 cell lysis buffer, RNase inhibitor, oligo-dT primer, and dNTP’s. Plates were spun for 1 minute to collect the liquid and cell at the bottom of the wells and immediately frozen. The Smartseq2 protocol was performed by the Israel National Center for Personalized Medicine (G-INCPM) as previously described [25] with 22 amplification cycles. Altogether, 6 plates were processed, each containing 96 individual cells (576 cells total), with 3 plates containing Six2-high cells and 3 plates containing Six2-low cells. Each one of the 6 plates was separately sequenced 1 × 50 bases on the Illumina HiSeq 2000 platform in the Israel National Center for Personalized Medicine (G-INCPM).

### Bulk RNA sequencing

We sorted Six2-high and Six2-low cells into separate 1.5 ml tubes, each containing RNA purification buffer. This was repeated in three experiments in which we used two types of RNA purification kits: the “single cell RNA Purification Kit” (51800, Norgen Biotek) was used in one experiment where we sorted 50,000 Six2-high and 50,000 Six2-low cells, and the Direct-zol RNA MiniPrep (R2050, Zymo Research) was used in two experiments where we sorted approximately 100,000 Six2-high cells and 400,000 Six2-low cells. Bulk total RNA was purified according to the manufacturer’s instructions and stored in −80°C. RNA was quantified on an Agilent BioAnalyzer (Agilent Technologies) and aliquots of 100-500ng were used to generate cDNA libraries using the TruSeq mRNA-Seq library kit (Illumina). Altogether, 6 libraries (3 replicates, each containing total RNA from Six2-high and Six2-low cells) were sequenced paired-end 2 × 125 bases on an Illumina HiSeq 2000 platform in the Israel National Center for Personalized Medicine (G-INCPM).

### Single cell RNAseq data preprocessing and gene expression analysis

Raw reads from 576 cells (6 × 96-well plates) were aligned by TopHat2 [58] to the mouse mm10 genome. Aligned reads were counted by HTSeq [59]. Data normalization and estimation of size factors was done by DESeq2 [60], resulting in a matrix of normalized gene expression counts.

We then filtered out 11 cells that expressed zero levels of the “housekeeping genes” Gapdh or Actb, resulting in 565 cells. We chose highly variable genes using the method by Macosko et al. [9] to select for genes whose variance exceeds those of other genes having a similar mean expression value (Fig. S2A). This step resulted in 647 highly variable genes, to which we added a list of genes from the literature that were previously shown to be involved in kidney development (Table S1). We also added an additional list of 48 genes from a previous single-cell qPCR study that we previously conducted on human fetal kidney cells [61] (Table S1), which, in retrospect, were not crucial to the identification of the different cell populations. These steps resulted in a gene expression matrix of 677 genes × 565 cells. Each gene was then modified-log-transformed [log2(1+expression)] and standardized by subtracting the mean, dividing by the standard deviation, and truncating to the range [−1,1].

We used tSNE [62] to project the 565 single cells profiles into a 2-dimensional plane (Fig. S2E) and used genes that are known to mark different populations in the fetal kidney to identify the various populations, including a population of 21 low quality cells that appears as a “mixture” of many cell types. This population of cells displayed low expression levels of the “housekeeping genes” Actb and Gapdh, as well as low DESeq size factors (Fig. S3). After removing these 21 low quality cells, the process of selecting for highly variable genes (and adding known genes related to kidney development as described above) was then repeated for the remaining high quality cells, resulting in a matrix of 728 genes × 544 cells, whose analysis is shown below. We note that once a sufficient number of highly variable genes are included, the ability to identify the different cell populations (Fig. 2A) is not very sensitive to the exact choice of genes.

### Creation of “bulk” *in-silico* transcriptomes representing the different cell populations

After identifying the different populations according to known gene markers from the literature (e.g. [6,7]), we created “bulk” *in-silico* transcriptomes representing the different cell populations by merging reads (*.bam files) from all cells belonging to each population (samtools merge). Again, aligned reads were counted by HTSeq [22] and data normalization and estimation of size factors was done by DESeq2 [23].

### Splicing analysis: Identification of splicing events and putative splicing regulators

rMATS [30] was used to detect splice isoform switching events between the “bulk” *in-silico* transcriptomes representing the different cell populations. In order to compare the mesenchymal populations (UM, CM) to the epithelial populations (PROX_1, PROX_2, LOH, DIST_CD), we performed the following comparisons: UM-PROX_1, UM-PROX_2, UM-LOH, UM-DIST_CD, CM-PROX_1, CM-PROX_2, CM-LOH, CM-DIST_CD. Additionally, we compared the mesenchymal populations (UM, CM) to the podocytes (PODO): UM-PODO and CM-PODO; See Table S2.

Selected splicing events (e.g. cassette exons) were visualized and validated using IGV [63] and Sashimi plots [64]. Gene Ontology (GO) enrichment of genes containing differential splicing events was done with ToppGene (https://toppgene.cchmc.org) [65]. rMAPS (http://rmaps.cecsresearch.org/) [51] was used to test for enrichment of binding motifs of RNA binding proteins (RBP’s) in the vicinity of alternatively spliced cassette exons in order to identify putative splicing regulators. A list of 84 RNA binding proteins (RBP’s) was obtained from the rMAPS website (http://rmaps.cecsresearch.org/Help/RNABindingProtein) [51–53]. Apart from the RNA binding motifs that are tested by the default settings in the rMATS website, we also tested additional UGG-enriched motifs that were previously found to be binding sites for the RNA binding proteins Esrp1 [19,31] and Esrp2 [38] (Table S4). For the RNA binding proteins Rbfox1 and Rbfox2, following [12] and the CISBP-RNA database [52] (http://cisbp-rna.ccbr.utoronto.ca) we assumed that both proteins (Rbfox1 and Rbfox2) preferentially bind to the same motif ([AT]GCATG[AC]) on mRNA.

## AUTHOR CONTRIBUTIONS

Study initiation and conception - B.D., A.U., and T.K.; Fetal kidney collection and dissociation - T.H.B.L, A.R., N.B.H., L.A., E.B., A.U., and T.K.; FACS sorting - D.I.; C1 experiments (data not used) - T.H.B.L.; Biomark experiments (data not used) - S.O.; Bulk RNA extraction - A.R., L.A., and A.U.; Bulk RNA quality checks – T.H.B.L.; Smartseq2 and RNA sequencing – S.G. and S.B.; Bulk and single cell RNA sequence preprocessing – N.B.H.; Single cell gene expression and splicing analysis – Y.W., N.B.H., and T.K.; Other intellectual contribution – Y.Y., N.P.S., and P.H.; Manuscript writing – Y.W. and T.K.

## ACKNOWLEDGEMENTS

We wish to thank Nelly Komorovsky for assistance in fetal kidney collection. Likewise, we wish to thank Jordan Kreidberg, Steve Potter, Oded Volovelsky, Morris Nehama, Tal Shay, Rotem Karni, and all members of our labs for useful comments and suggestions.

## FUNDING

Y.W., T.H.B.L., N.B.H., S.O., E.B., Y.Y., and T.K., were supported by the Israel Science Foundation (ICORE no. 1902/12 and Grants no. 1634/13 and 2017/13), the Israel Cancer Association (Grant no. 20150911), the Israel Ministry of Health (Grant no. 3-10146), and the EU-FP7 (Marie Curie International Reintegration Grant no. 618592). The funders had no role in study design, data collection and analysis, decision to publish, or preparation of the manuscript.

## COMPETING INTERESTS

The authors have declared that no competing interests exist.

## SUPPORTING INFORMATION LEGENDS

**Supplementary information:** Supplementary figures.

**Table S1:** A list of genes from the literature that were previously shown to be involved in kidney development and an additional list of 48 genes from a previous single-cell qPCR study that we previously conducted on human fetal kidney cells [61]. We note that once a sufficient number of genes are included, the ability to identify the different populations (Figs. 1B, 2A) is not very sensitive to the exact choice of genes.

**Table S2:** rMATS tables for cassette exons that are alternatively spliced between the different populations.

**Table S3:** Gene Ontology (GO) enrichment analysis results from ToppGene.

**Table S4:** A list of RNA motifs used for identifying putative splicing regulators.

**Table S5:** Single-cell gene expression values.

**Table S6:** Lists of cells in each population.

**Program:** A compressed directory containing a short matlab program and datasets for single-cell data visualization.

