## Supplementary Information and figures for "Single-cell RNA sequencing reveals mRNA splice isoform switching during kidney development"

[1]*Department of Bioengineering and Bar-Ilan Institute of Nanotechnology and Advanced Materials (BINA), Bar-Ilan University, Ramat Gan, Israel 52900*
[2]*The Mina and Everard Goodman Faculty of Life Sciences, Bar-Ilan University, Ramat-Gan, Israel 52900*
[3]*The Nancy and Stephen Grand Israel National Center for Personalized Medicine (G-INCPM), Weizmann Institute of Science, Rehovot, Israel 7610001*
[4]*Pediatric Stem Cell Research Institute, Edmond and Lily Safra Children's Hospital, Sheba Medical Center, Tel-Hashomer, Israel 52621*
[5]*Division of Pediatric Nephrology, Sheba Medical Center, Tel-Hashomer, Israel 52621*
[6]*Sackler Faculty of Medicine, Tel-Aviv University, Tel-Aviv, Israel*
[7]*Dept. Human Genetics, Leiden University Medical Center, Leiden, The Netherlands*
[*]*Co-first authors*
[**]*Co-senior authors*

[8]**Correspondence to:**
Tomer Kalisky
Department of Bioengineering, Bar-Ilan University, Ramat Gan, Israel 52900


**Calculation of the likelihood function for inclusion levels of the KTS+/- isoforms of the gene Wt1 (Fig. S20B):**

To calculate the likelihood function for the KTS inclusion level $\psi$, which is defined as the fraction of transcripts that include the KTS segment out of the total number of transcripts that either include the KTS segment or skip over it [1], we counted the number of reads that span the junction between exons 9 and 10, while either including the KTS segment ($I$) or skipping it ($S$) (Fig. S20).

Since:

$$I|\psi{\sim}Binomial(N = I + S, p = \psi)$$

$$Prob(I|N = I + S, p = \psi) = \binom{I + S}{I} \psi^I (1 - \psi)^S$$

Note that since we only counted reads that span the junction between exons 9 and 10, the effective lengths [1] (that is, the number of unique isoform-specific read positions) of the KTS-inclusion isoform and the KTS-skipping isoform are taken to be the same.

Therefore, for given values of $I$ and $S$, we get the likelihood:

$$L(\psi|I, S) = \binom{I + S}{I} \psi^I (1 - \psi)^S$$

Where the, maximum likelihood is $\psi_{ML} = I/(I + S)$.

**Supplementary figures**

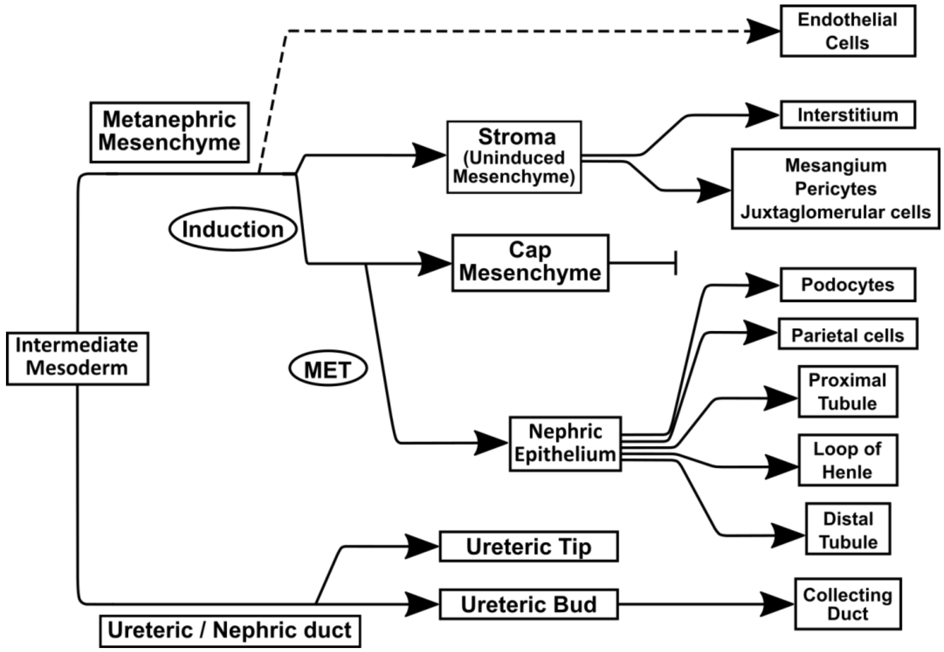


**Figure S1:** A sketch of the various cell lineages that co-exist in the nephrogenic zone of the developing fetal kidney [2].

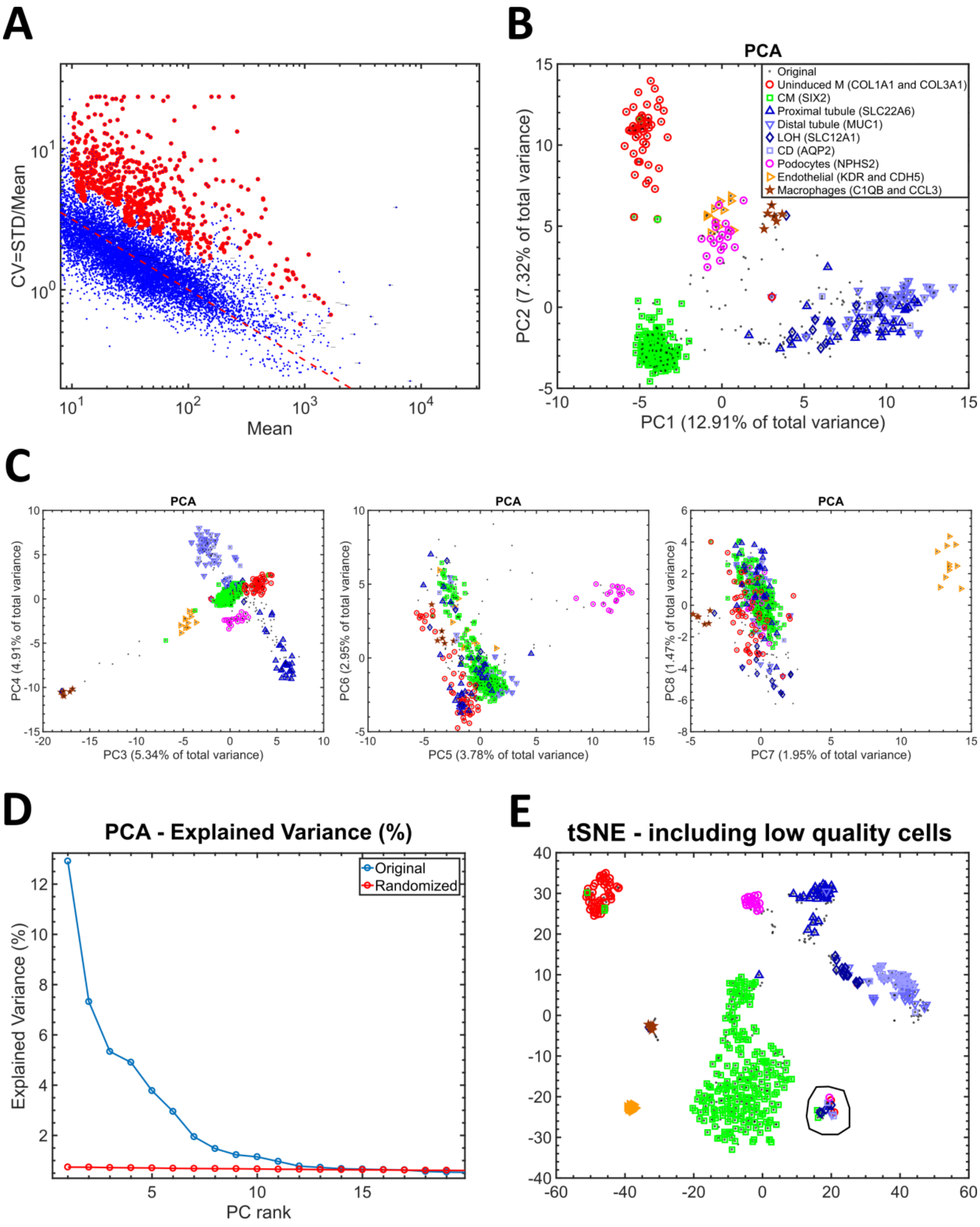


**Figure S2:** Preprocessing of single-cell data from the Smartseq2 protocol and removing low quality cells. Raw reads from 576 cells (6 x 96-well plates) were aligned to the mouse mm10 genome and the numbers of reads that align to each gene were counted and normalized. We filtered out 11 cells that expressed zero levels of the “housekeeping genes” Gapdh or Actb, resulting in 565 cells. (A) We chose highly variable genes. We followed the method by Macosko et al. [3] to select for genes whose cell-to cell variance exceeds that which would have resulted from a Poisson distribution. These are actually genes whose cell to cell heterogeneity cannot be attributed to random distribution of transcripts between wells and are therefore likely to be actively over-expressed or under-expressed in the different cell populations. We first removed all non-expressed genes and plotted the mean vs. coefficient of variance (CV=STD/Mean) for each gene (blue dots). Then, we filtered out genes whose normalized counts are less than 10. We divided the mean expression (horizontal axis) into 100 equally sized bins in log10 space, and for each bin we calculated the mean and dispersion (=standard deviation) of the CV (vertical axis). We chose only genes whose CV exceeded the mean CV within their respective bin by at least one standard deviation (red large dots). This actually chooses genes whose variance exceeds those of other genes having a similar mean expression value. This step resulted in 647 highly variable genes, to which we added a list of genes from the literature that were previously shown to be involved in kidney development (Table S1). We also added an additional list of 48 genes from a previous single-cell qPCR study that we previously conducted on human fetal kidney cells [4] (Table S1), which, in retrospect, were not crucial to the identification of the different cell populations. These steps resulted in a gene expression matrix of 677 genes x 565 cells. Each gene was then modified-log-transformed [log2(1+expression)] and standardized by subtracting the mean, dividing by the standard deviation, and truncating to the range [-1,1] (B) Principal Components Analysis (PCA) plot of the resulting matrix consisting of 677 genes vs. 565 cells. The first two principal components (PC's 1-2) show a triangular shaped structure whose vertices correspond to the un-induced mesenchyme (UM), cap mesenchyme (CM), and epithelial populations. (C) Higher principal components (PC's 3-4, 5-6, etc.) capture additional heterogeneity arising from other cell populations such as macrophages, podocytes, and endothelial cells. (D) The explained variance of each principal component. PC's 14 and onwards do not explain more variance than is explained by a randomized matrix, that is, a matrix in which each gene (=row) was randomly permuted. (E) A tSNE plot of 565 single cell profiles, each consisting of 677 highly variable genes. Cells over-expressing genes that were previously shown to mark different populations are marked by additional symbols (as in Fig. 2A in the main text). We identified a population of 21 low quality cells that appears as "mixture" of many cell types (circled). These 21 low quality cells were filtered out, resulting in a 544 cells. The process of selecting for highly variable genes (and adding known genes related to kidney development as described above) was then repeated for the remaining high quality cells, resulting in a matrix of 728 genes x 544 cells, whose analysis is shown in the main text.

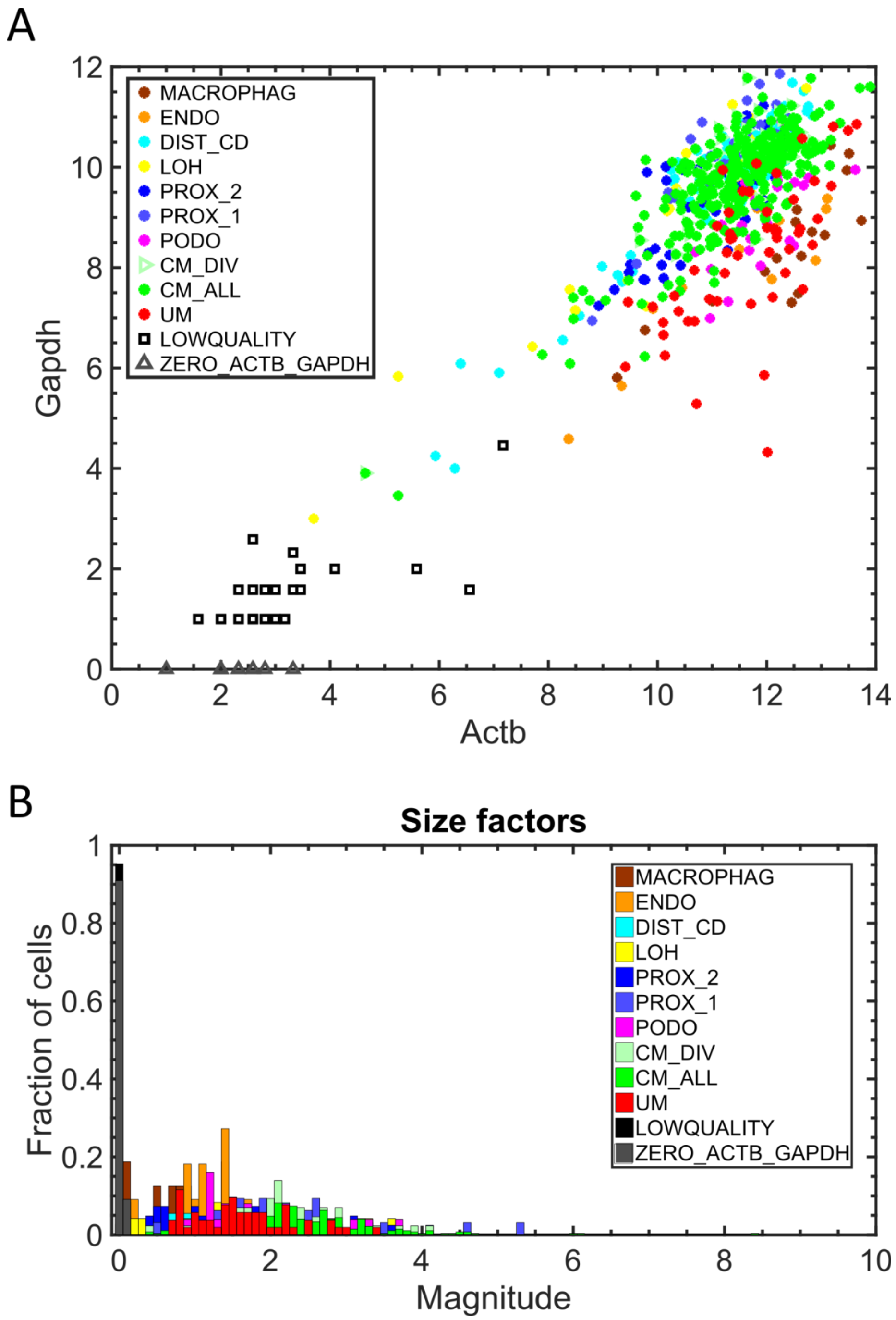


**Figure S3:** Removing low quality cells. (A) The population of 21 low quality cells (black squares) have low expression levels of the "housekeeping genes" Actb and Gapdh. (B) Likewise, they have low DESeq size factors [5,6]. The annotations "CM_ALL" and "CM" are used interchangeably to represent all cells that were classified as belonging to the cap mesenchyme. "CM_DIV" represents a subset of cells from the cap mesenchyme that are presumably dividing since they over-express the genes Mki67 and Top2a that are known to be over-expressed during the S-G2-M phase of the cell cycle.

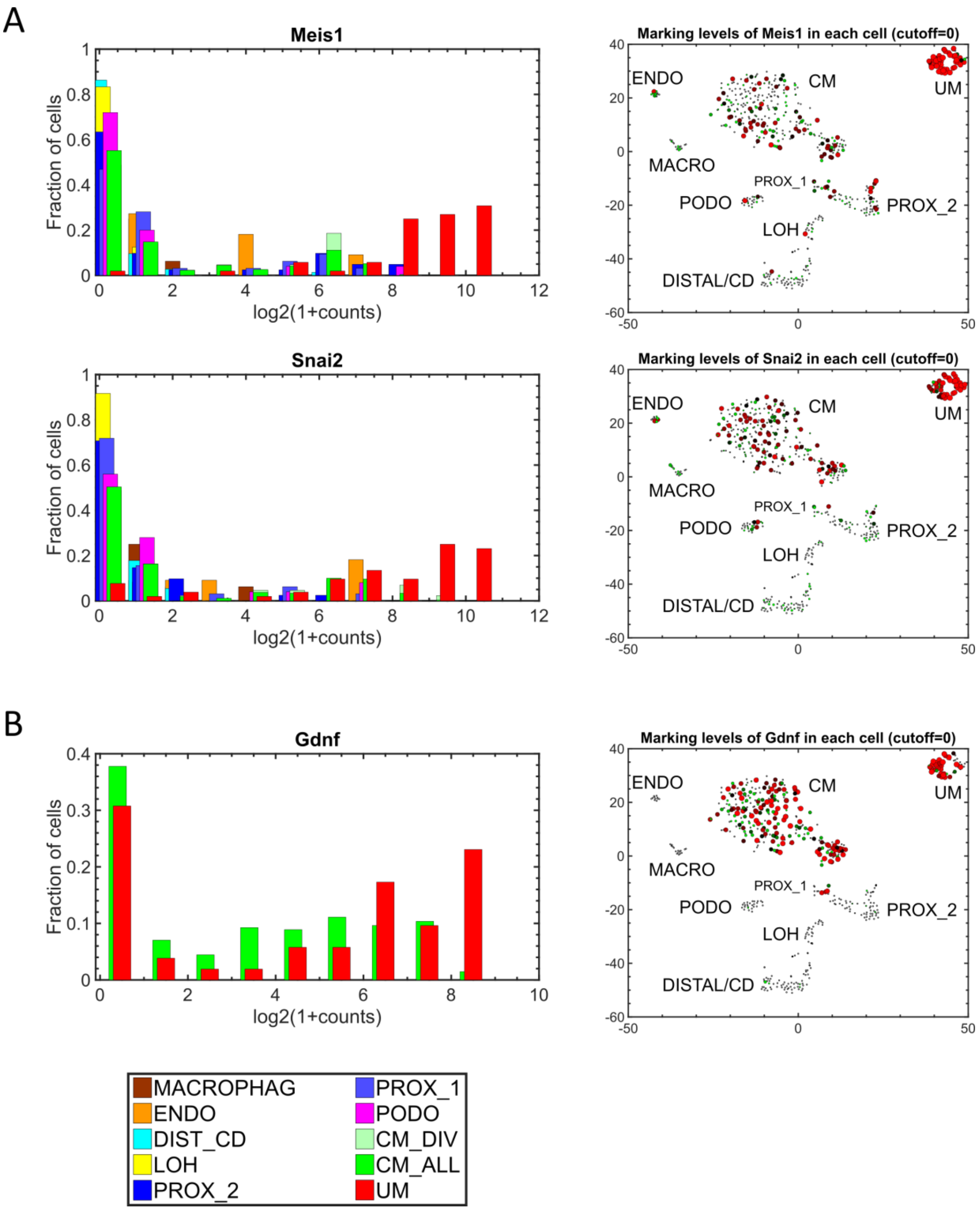


**Figure S4:** Confirmation of cell population identity. Shown are histograms and tSNE plots marking genes that are known to be over-expressed in the un-induced mesenchyme (UM) and stroma. Meis1 [7,8] and Snai2 [9] are more specific to the UM while Gdnf [8,10,11] is expressed in the un-induced mesenchyme (UM) and cap mesenchyme (CM) as previously shown [8].

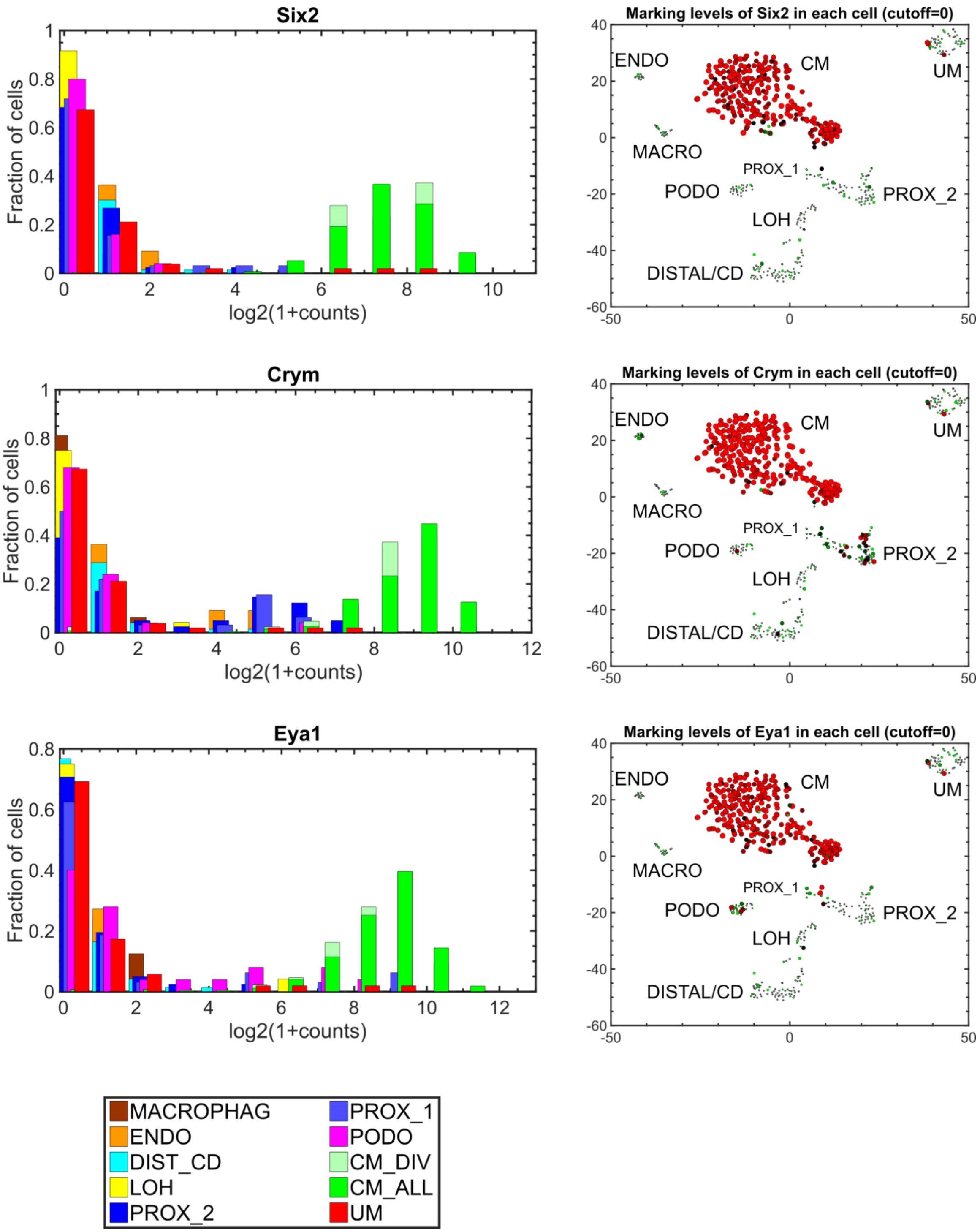


**Figure S5:** Confirmation of cell population identity. Shown are histograms and tSNE plots marking genes that are known to be over-expressed in the cap mesenchyme (CM) [11,12]. Note that Crym is also moderately expressed in the early epithelial structures (PROX_1) and proximal tubules (PROX_2) [8,10].

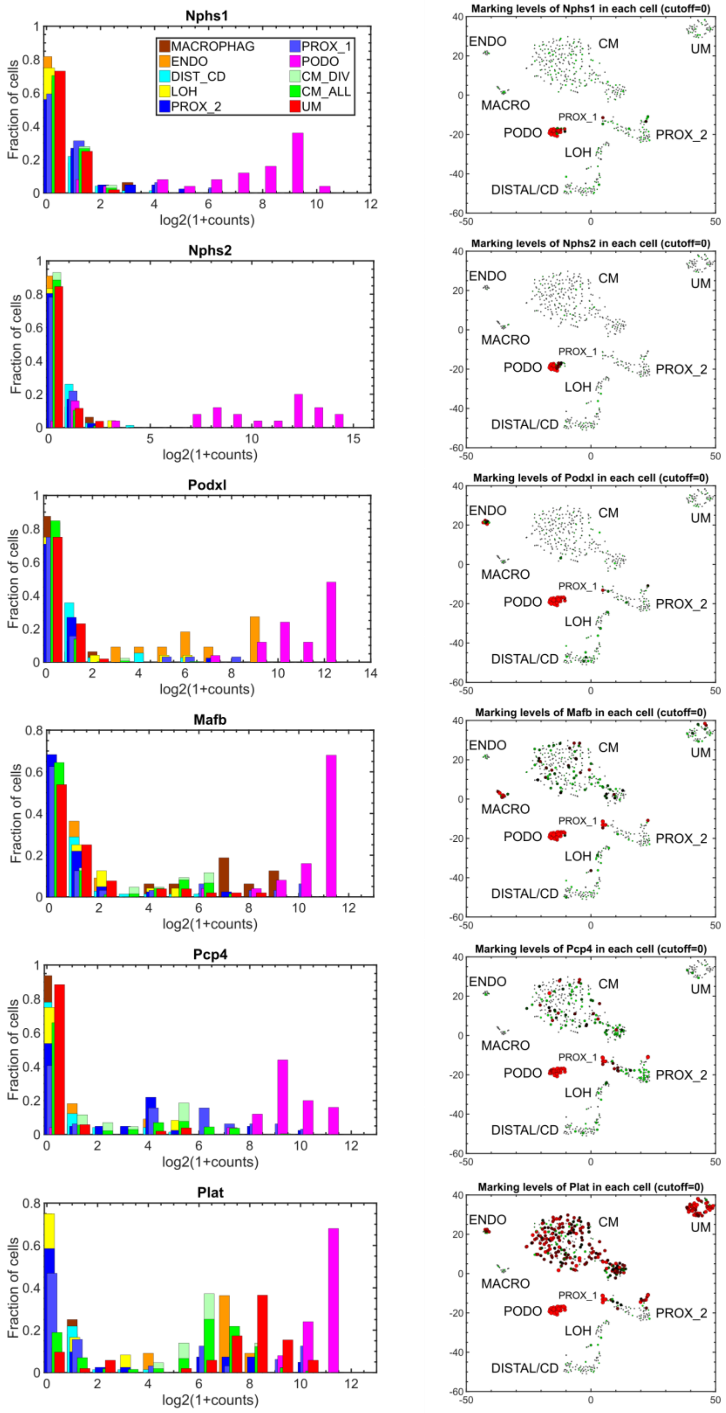


**Figure S6:** Confirmation of cell population identity. Shown are histograms and tSNE plots marking genes that are known to be over-expressed in the podocytes (PODO) [11]. Note that Plat is also moderately expressed in other early developmental lineages (CM, UM, PROX_1) [8,10].

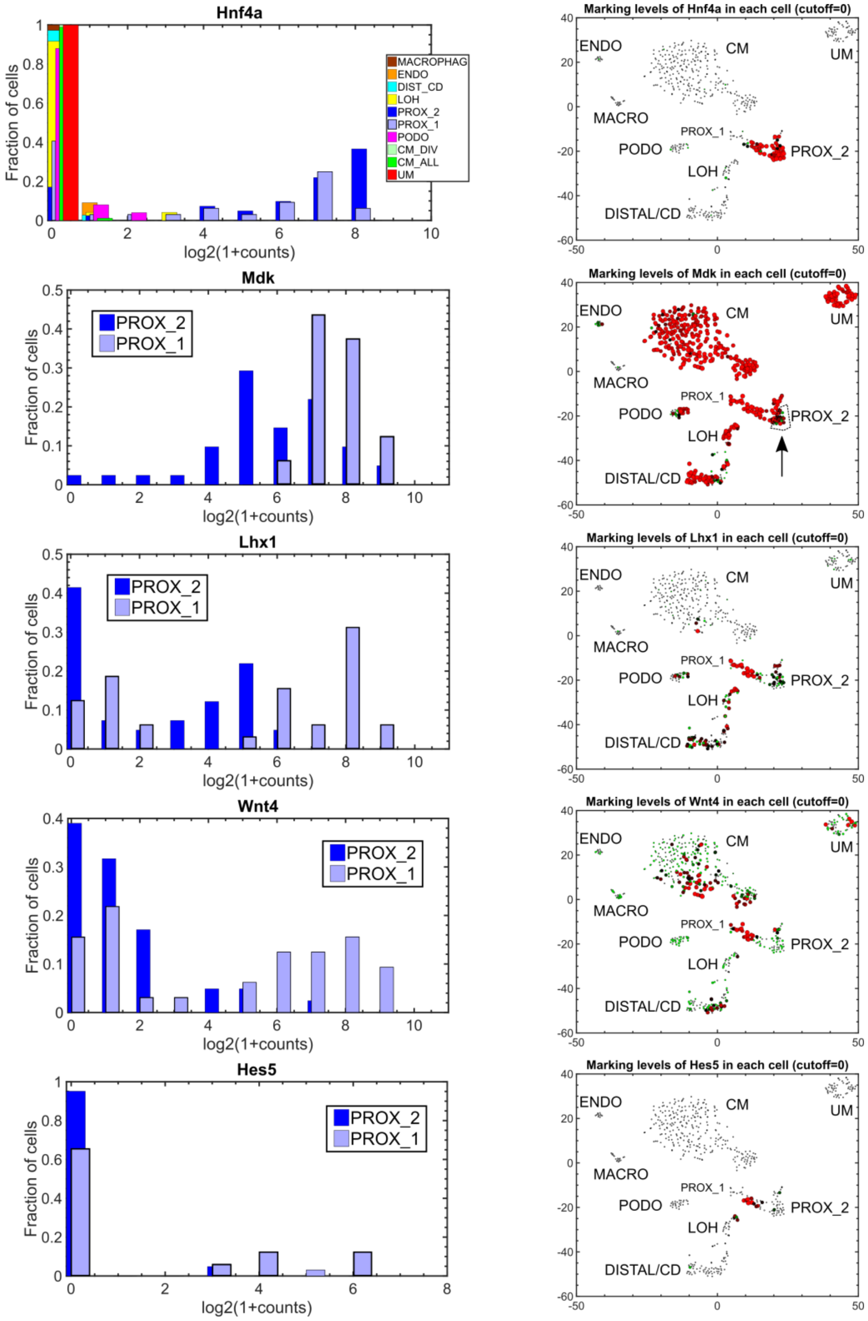


**Figure S7:** Confirmation of cell population identity. Shown are histograms and tSNE plots marking genes that are known to be over-expressed in the early epithelial structures (PROX_1) and proximal tubules (PROX_2) or genes that distinguish between them. Both populations express high levels of the proximal tubular marker Cdh6 [11,13] (Fig. 2D in main text). Hnf4a [7] is expressed in both PROX_1 and PROX_2, while Mdk [7,8], Lhx1 [7,11,14], Wnt4 [11,15,16], and Hes5 [7] are higher in the early epithelial structures (PROX_1) – which are presumably the pre-tubular aggregates, renal vesicles, and C/S-shaped bodies [8,10].

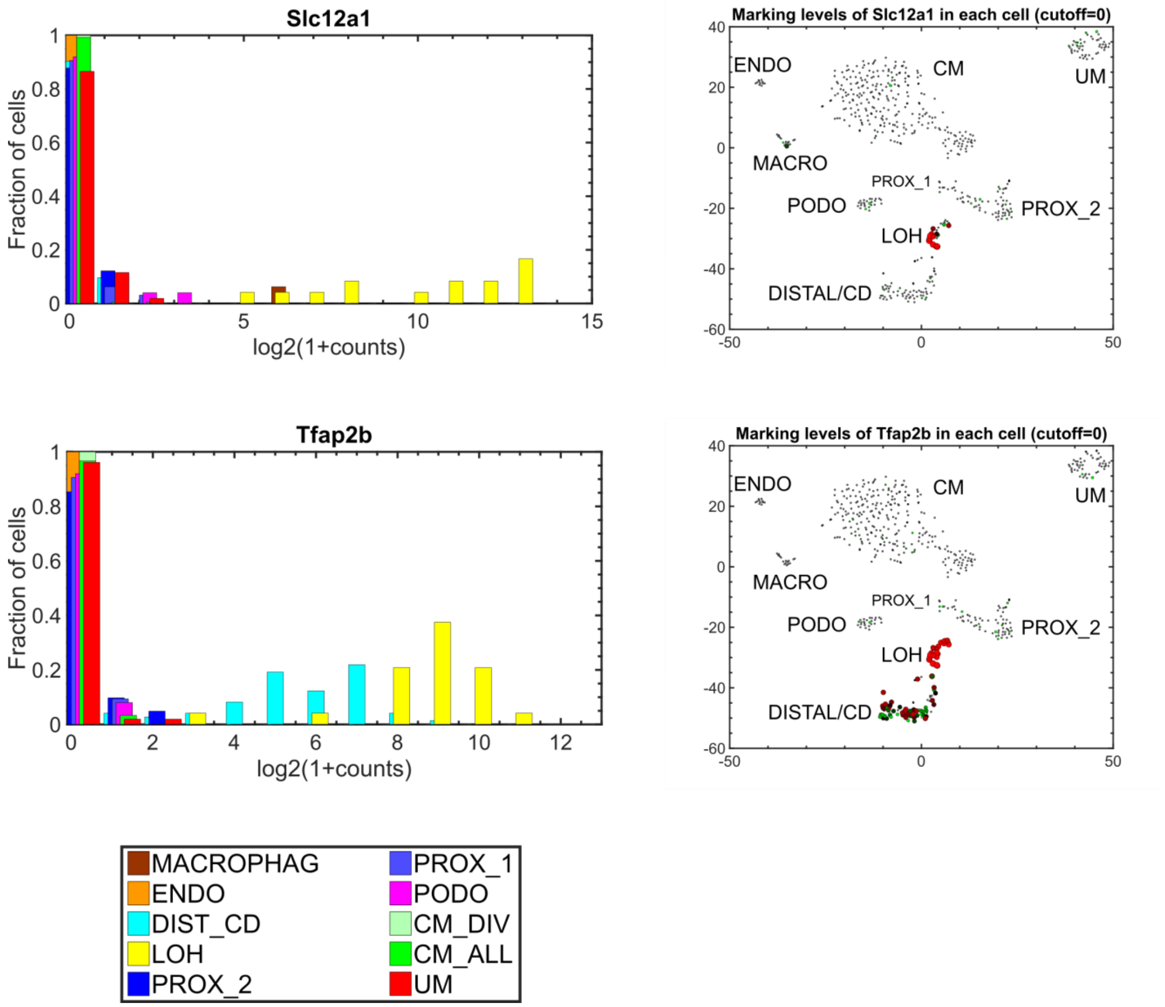


**Figure S8:** Confirmation of cell population identity. Shown are histograms and tSNE plots marking genes that are known to be over-expressed in the loop of Henle (LOH). Note that Tfap2b is also moderately expressed in the distal tubule and collecting duct (DIST_CD) [8,10].

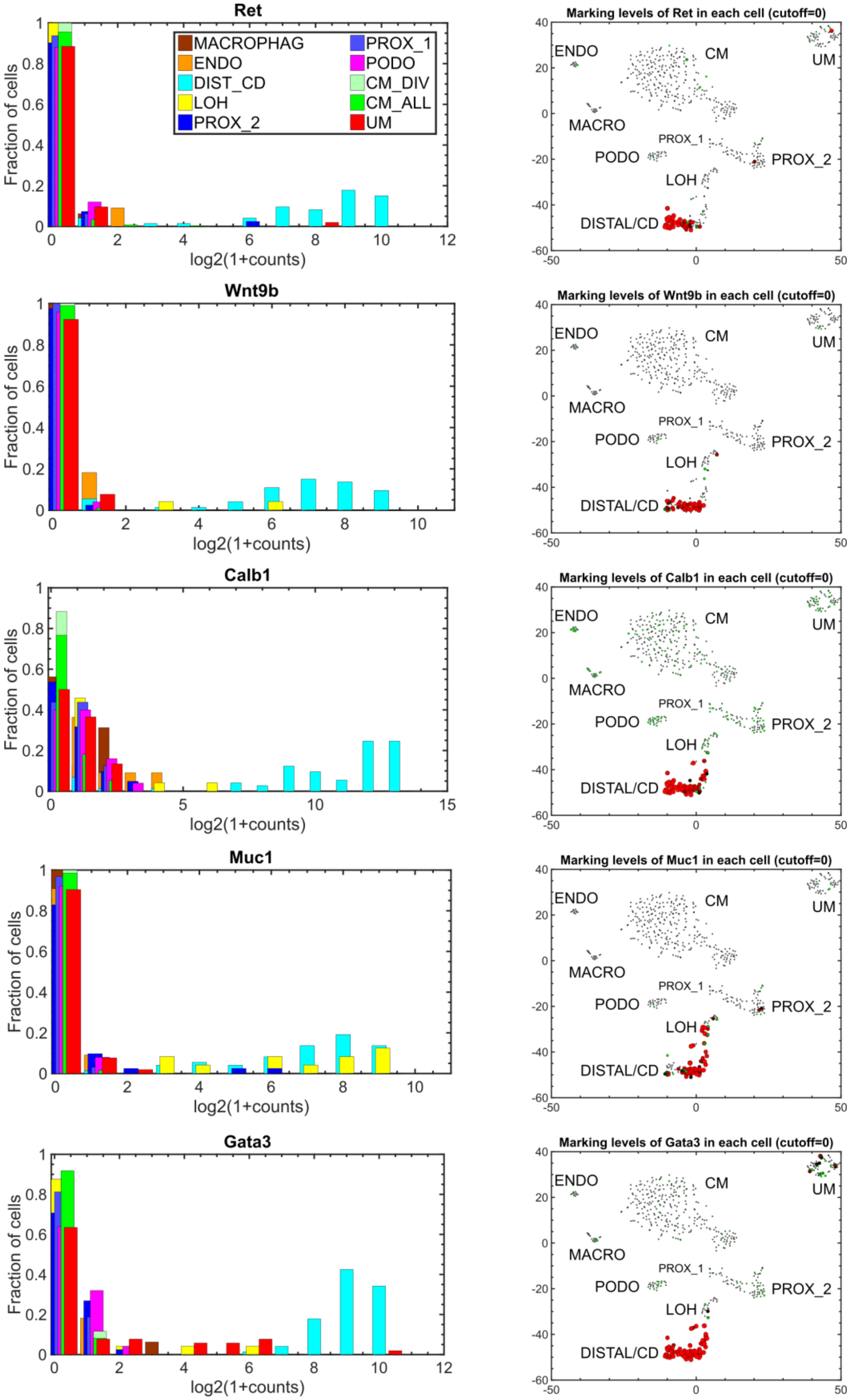


**Figure S9:** Confirmation of cell population identity. Shown are histograms and tSNE plots marking genes that are known to be over-expressed in the distal tubule and/or collecting duct (DIST_CD) [8,11,16–18]. We found it difficult to distinguish between the two populations, probably due to the small number of cells in our analysis. Note that the genes Ret and Wnt9b, which are known to mark the collecting duct, are slightly more restricted that the other genes which are known to mark both the collecting duct and the distal tubules.

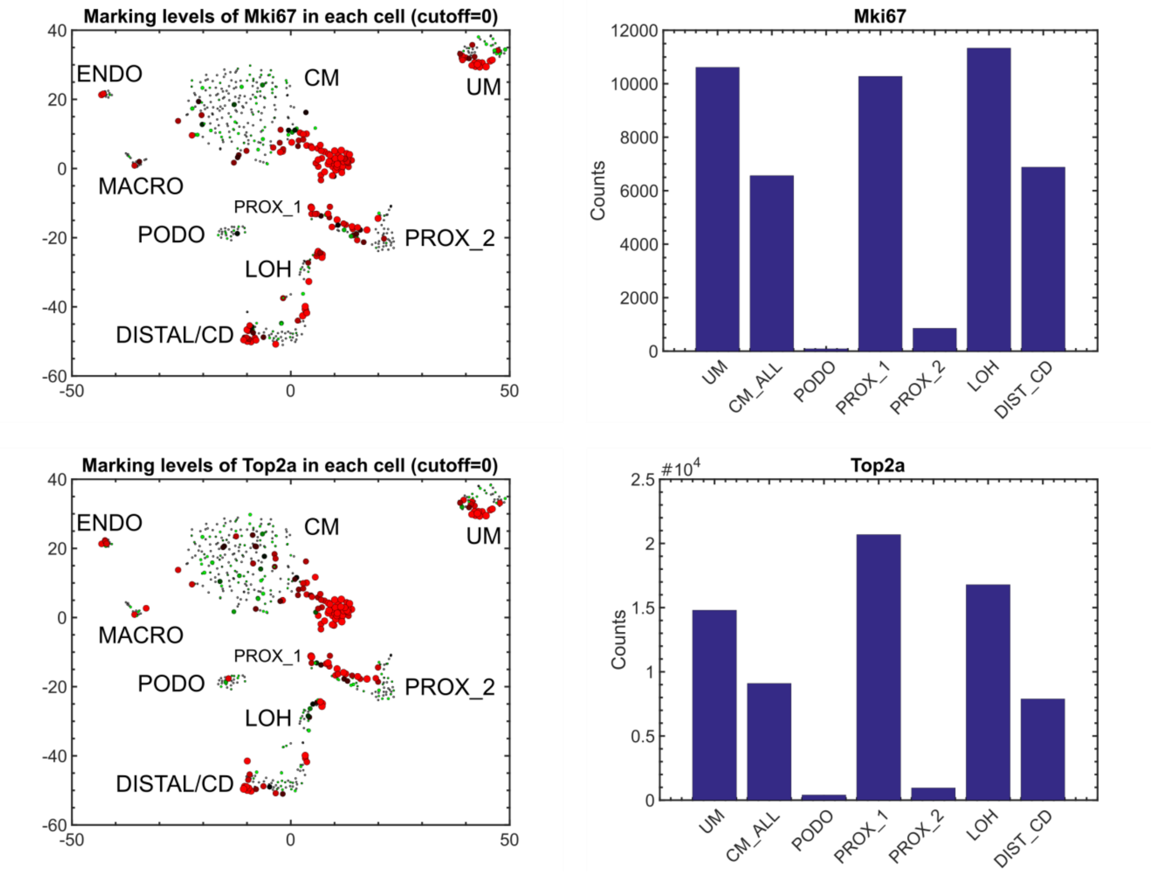


**Figure S10:** Cell division occurs in most cell populations of the developing kidney. Shown are tSNE plots marking the genes Mki67 and Top2a - genes that are typically expressed during the S-G2-M phases of the cell cycle - as well as barplots of their expression levels within the *in-silico* “bulk” transcriptomes representing the different populations. Note the higher expression in the early epithelial structures (PROX_1) with respect to the proximal tubules (PROX_2).

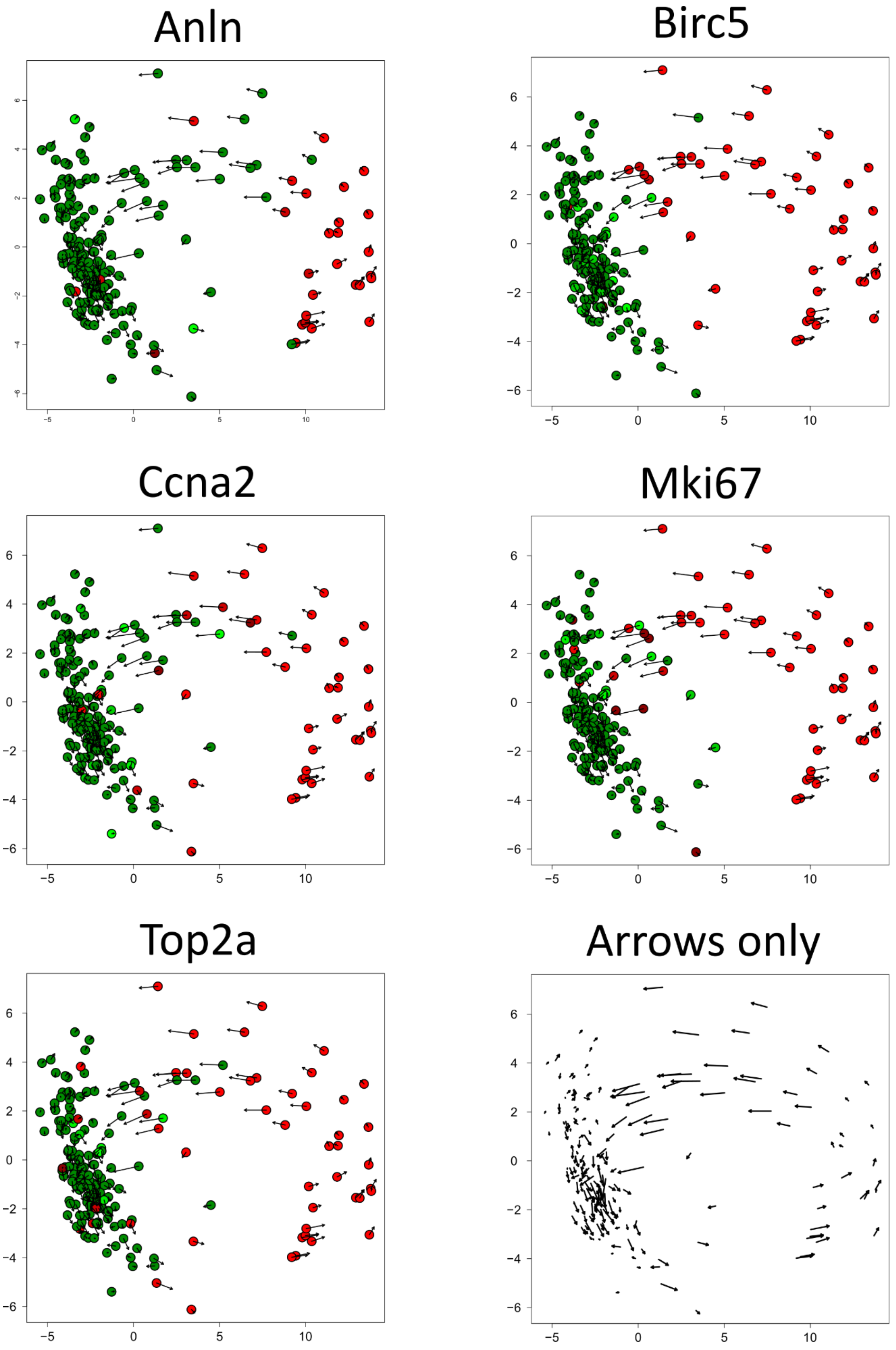
Anln
Birc5
Ccna2
Mki67
Top2a
Arrows only

**Figure S11:** RNA velocity [19] of all Six2-high cells shows a consistent directional "flow" along the circular manifold created by the cells of the cap mesenchyme (CM) in gene expression space (PC 1 vs. 2). Each cell is represented by a dot. The arrows represent the directionality of each cell in gene expression space that is inferred from the difference between the spliced transcriptome (=present state) vs. yet-unspliced transcriptome (=near future state) [19]. Circle fill colors represent expression levels of selected genes (Red – high expression, green – low expression). It can be seen that cells over-expressing genes such as Top2a and Mki67 - genes that are known to be over-expressed in the S-G2-M phases of the cell cycle - are located in a specific segment of the circular manifold representing the S-G2-M segment of the cell cycle. At this segment the arrows are longest, indicating a rapidly changing transcriptional state.

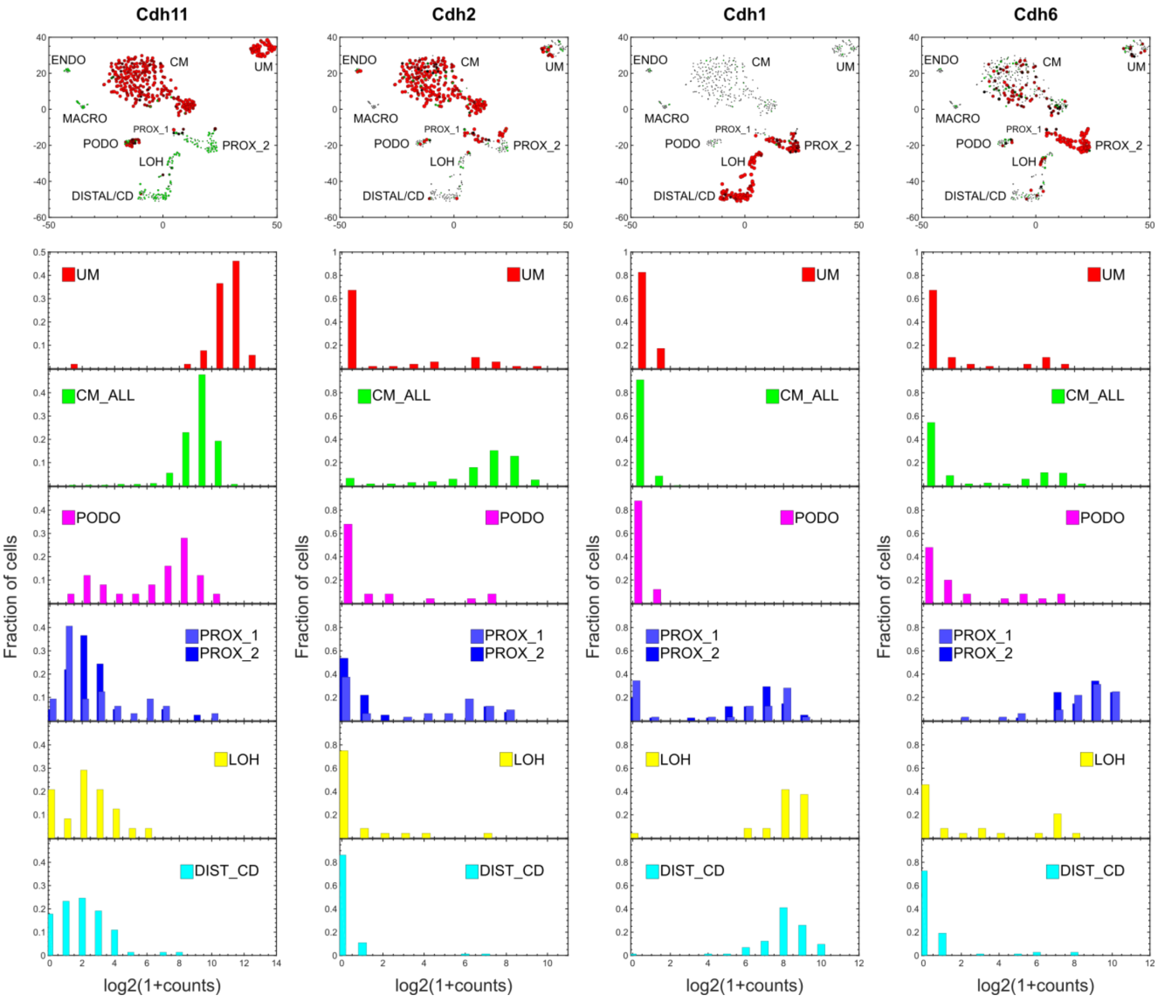


**Figure S12:** Mesenchymal to epithelial transition (MET) during kidney development. Shown are histograms and tSNE plots marking genes that are over-expressed in mesenchymal cells (Cdh11 [13] and Cdh2 [20]) and epithelial cells (Cdh1 and Cdh6 [13]). Note that Cdh11 is expressed in three levels: high expression in the un-induced mesenchyme (UM), moderate expression in the cap mesenchyme (CM), and low expression in the remaining epithelial populations (PROX_1, PROX_2, LOH, DIST_CD). In the podocytes we observed Cdh11 to have a bimodal distribution consisting of medium and low expression levels, indicating two distinct levels of differentiation.

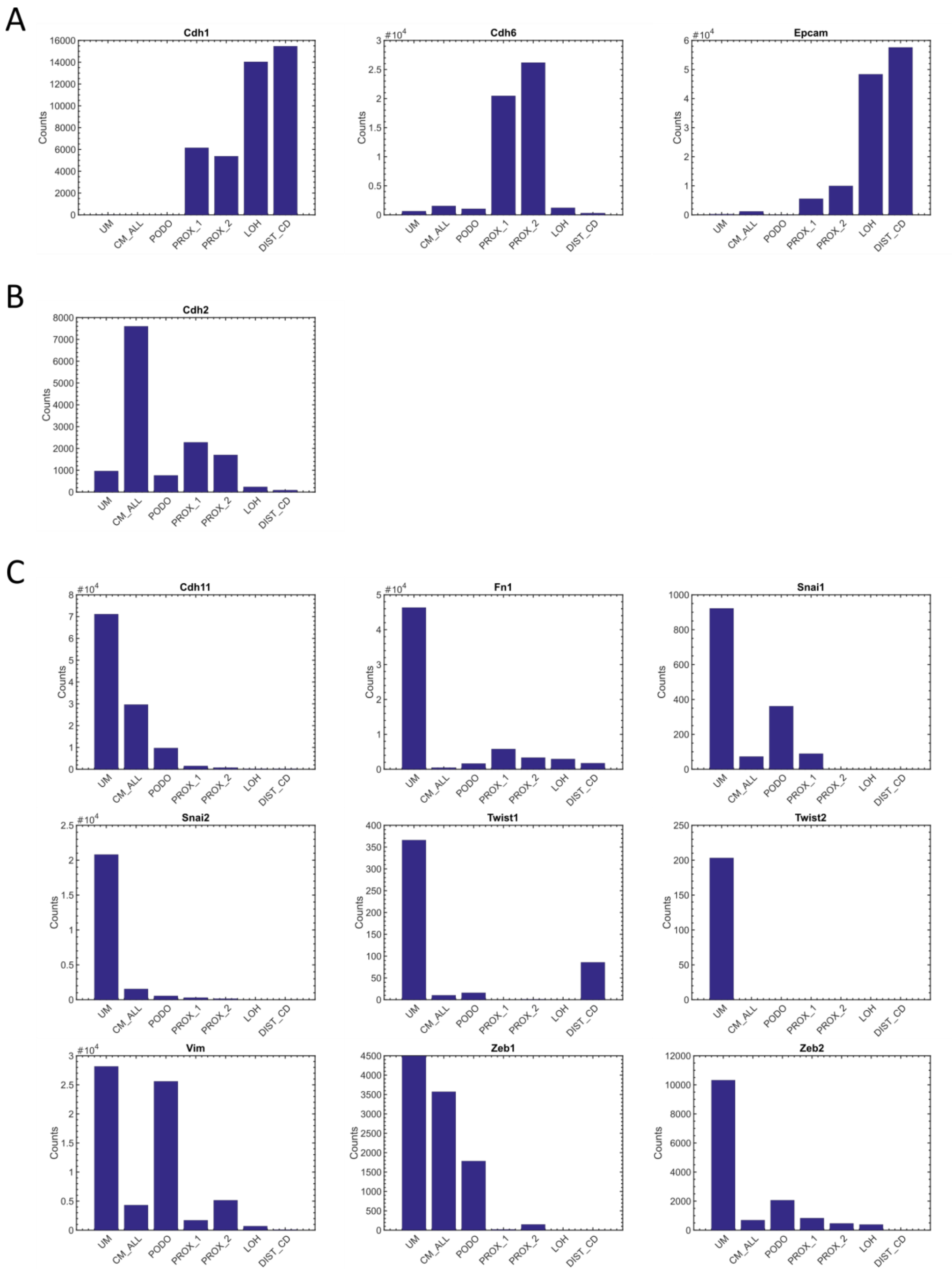


**Figure S13:** Mesenchymal to epithelial transition during kidney development. Shown are barplots of expression levels of selected genes within the *in-silico* “bulk” transcriptomes representing the different populations. (A) Epithelial markers (B) The mesenchymal marker Cdh2 [20] is over-expressed in the cap mesenchyme (CM) (C) Other mesenchymal markers. Note that Vimentin (Vim) is also highly expressed in the podocytes.

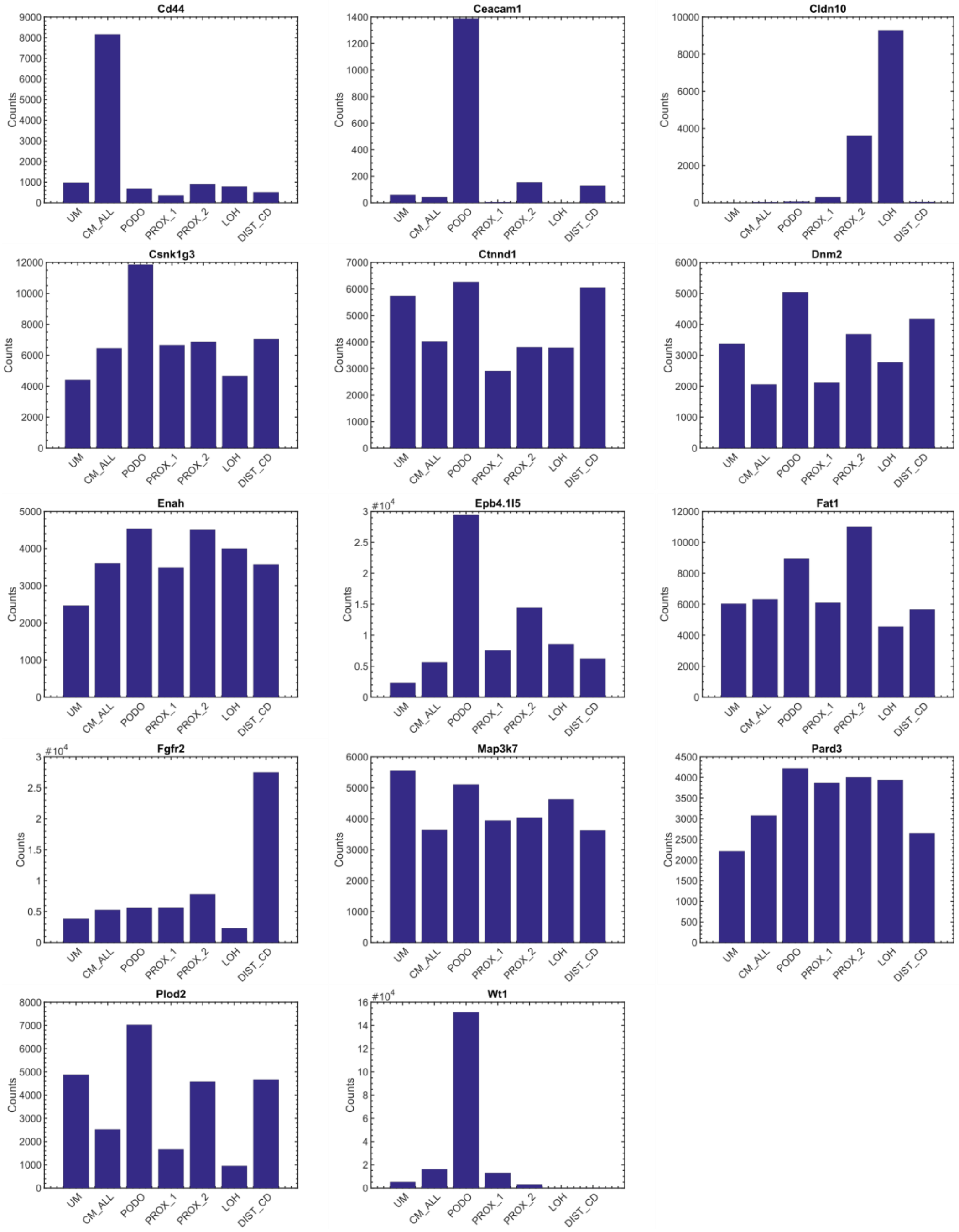


**Figure S14:** No obvious correlation between gene expression levels and inclusion levels of associated cassette exons. Shown are expression levels of selected genes in the *in-silico* “bulk” transcriptomes representing the different cell populations. For example, exons in the genes Dnm2 and Map3k7 have low inclusion levels in the mesenchymal populations and high inclusion levels in the epithelial populations (Fig. 3) but almost constant levels of gene expression.

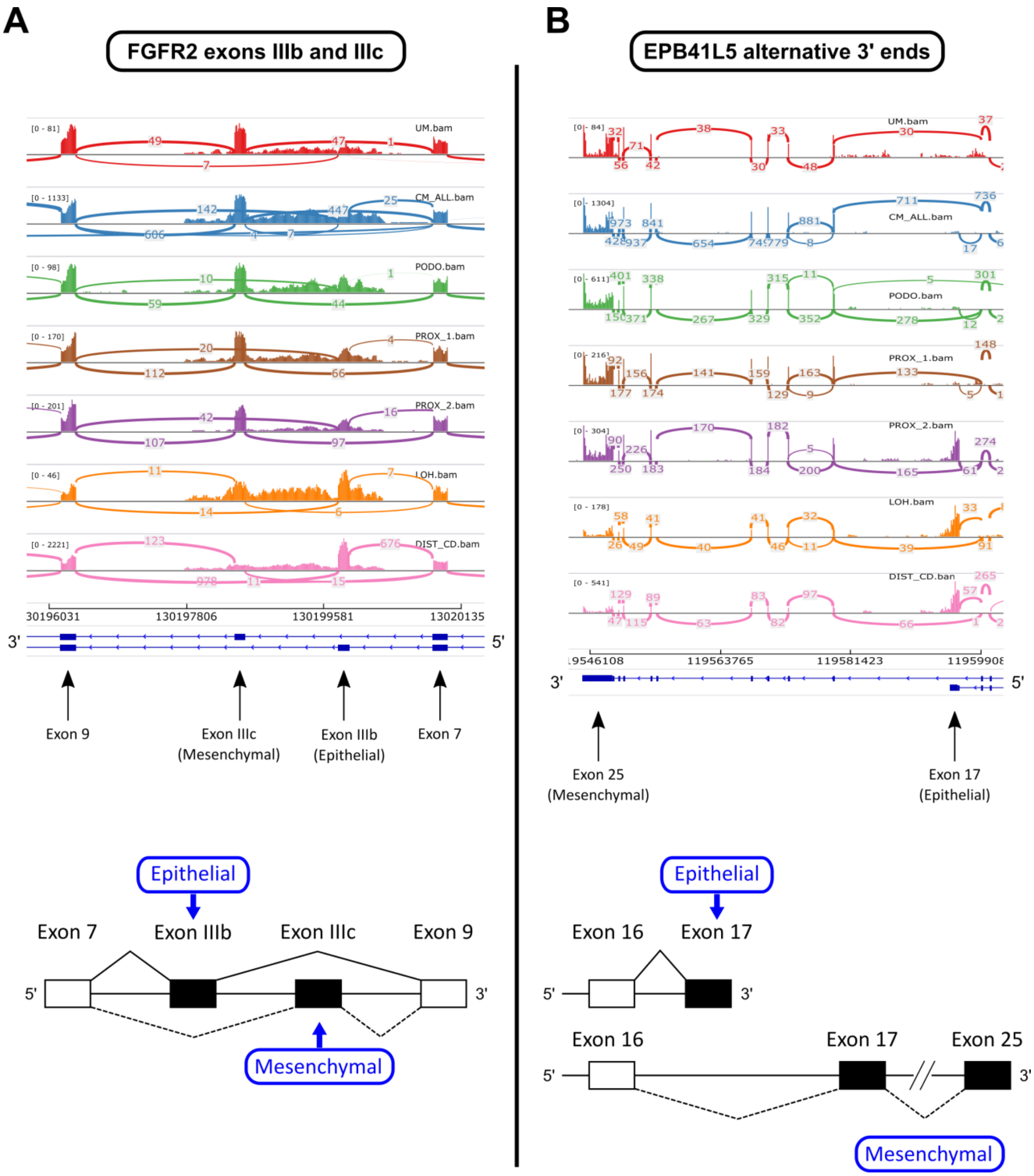


**Figure S15:** The genes Fgfr2 and Epb41l5 switch between mesenchymal and epithelial isoforms during kidney development. (A) The mesenchymal isoform of Fgfr2 (Fgfr2-IIIc) [21,22] is predominantly expressed in the mesenchymal and early developmental cell populations (UM, CM, PODO, and to some extent PROX_1 and PROX_2) while the epithelial isoform (Fgfr2-IIIb) is expressed mostly in the mature epithelial cell populations (LOH and DIST_CD). Note that the tubular epithelial cell populations PROX_1, PROX_2, and LOH contain varying mixtures of the two isoforms. The dominance of the mesenchymal isoform of Fgfr2 (Fgfr2-IIIc) in the un-induced mesenchyme (UM) and cap mesenchyme (CM) is in agreement with previous observations that deletion of Fgfr2-IIIc (along with conditional deletion of Fgfr1 in the metanephric mesenchyme) results in poorly formed metanephric mesenchyme and unbranched ureteric buds [23,24]. (B) The longer mesenchymal isoform of Epb41l5 [21] is predominantly expressed in the mesenchymal and early developmental cell populations (UM, CM, PODO, and PROX_1) while the shorter epithelial isoform is expressed mostly in the mature epithelial cell populations (LOH and DIST_CD). The proximal tubular cell population (PROX_2) contains a mixture of the two isoforms.

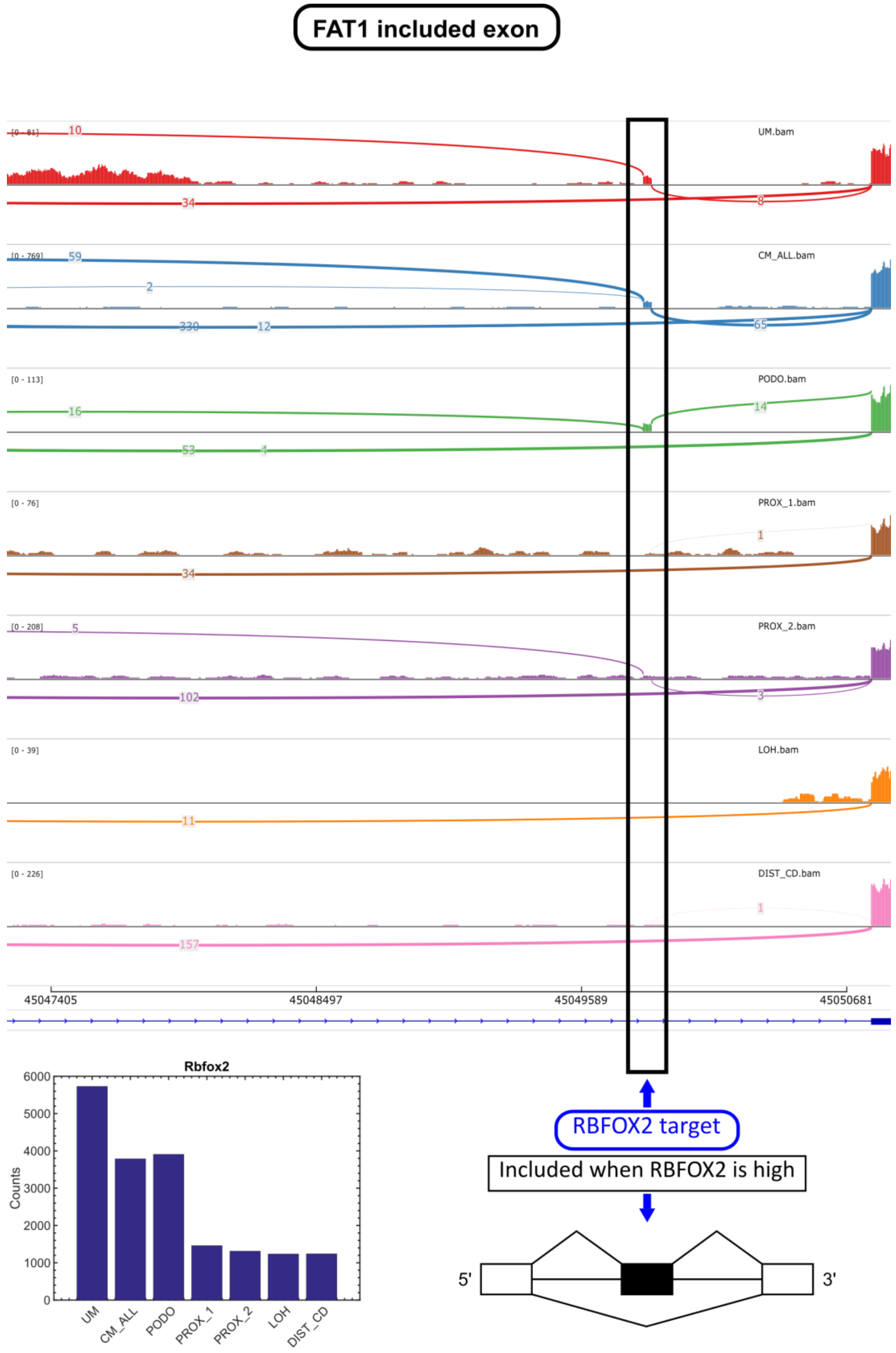


**Figure S16:** The gene Fat1, a known target of the splicing regulator Rbfox2 [25], undergoes splice isoform switching during kidney development. A cassette exon is expressed in the mesenchymal cell populations (UM, CM) and podocytes (PODO), and repressed in the epithelial cell populations (PROX_1, PROX_2, LOH, and DIST_CD), in accordance with the expression levels of Rbfox2. This indicates that Rbfox2 acts as a splicing regulator during kidney development.

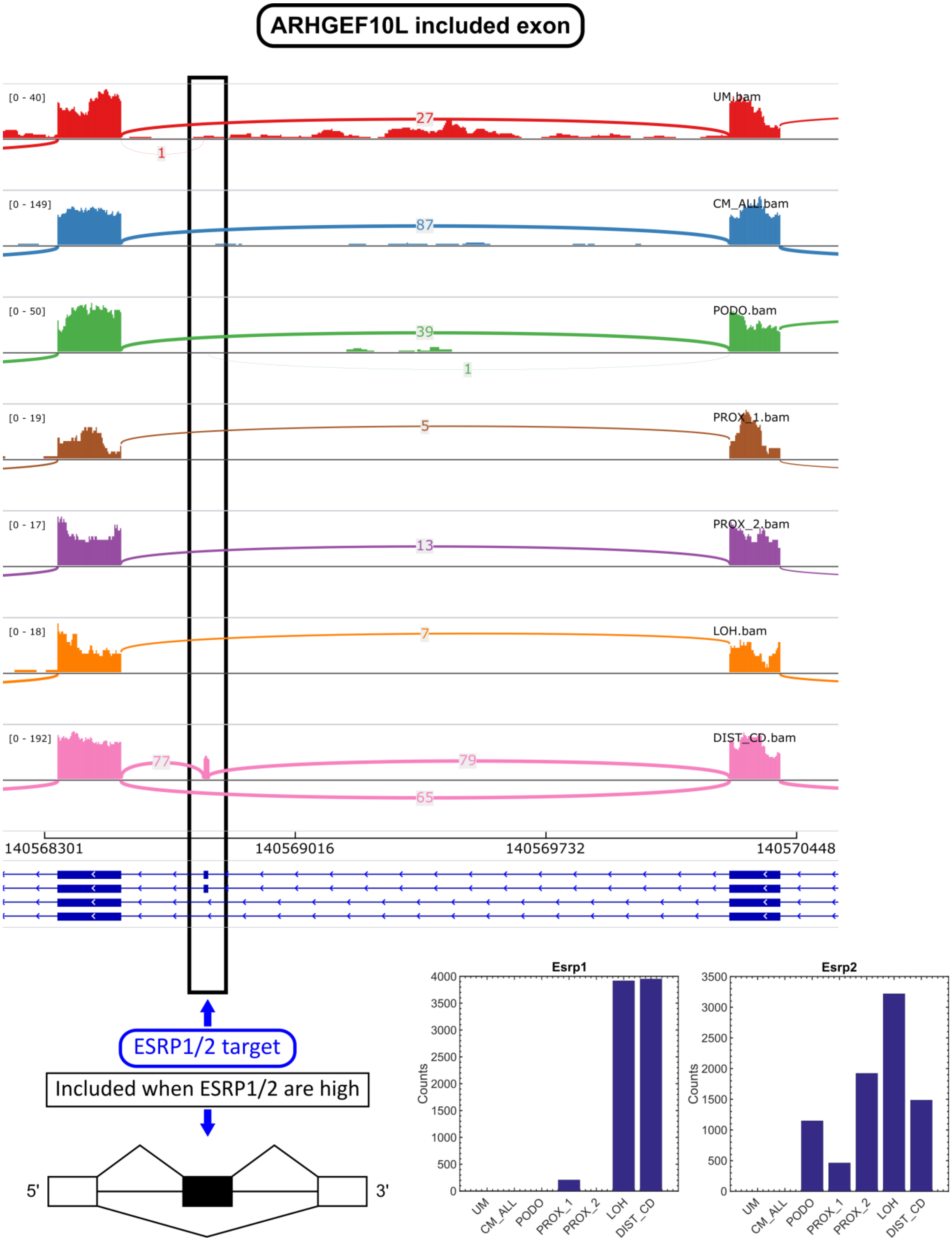


**Figure S17:** The gene Arhgef10l, a known target of the splicing regulators Esrp1/2 [26], undergoes splice isoform switching during kidney development. A cassette exon is repressed in the mesenchymal populations (UM,CM) and podocytes (PODO) and over-expressed in the distal tubules/collecting duct (DIST_CD). This is in accordance with the expression levels of Esrp1 and Esrp2, which are jointly under-expressed in the mesenchymal populations (UM,CM) and over-expressed in the epithelial populations (mainly LOH and DIST_CD). This indicates that Esrp1 and Esrp2 act as splicing regulators during kidney development. Note that Arhgef10l is sufficiently expressed only in the mesenchymal populations (UM, CM), podocytes (PODO), and distal tubules/collecting duct (DIST_CD) but not so much in the other epithelial populations (PROX_1, PROX_2, and LOH), thus limiting our ability to reliably measure the inclusion levels of the alternatively spliced exon in these populations.

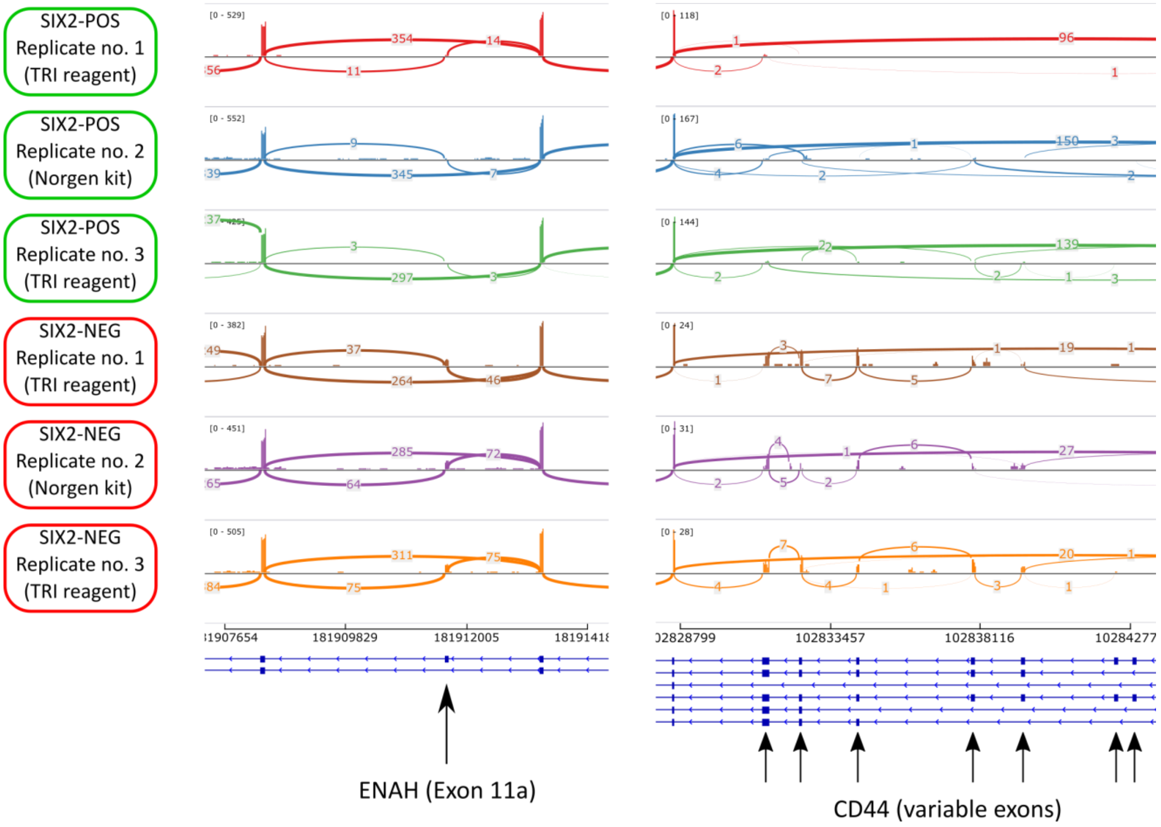


**Figure S18:** Alternative splicing in the genes Enah and CD44, two genes that are known to undergo splice isoform switching during EMT [21,22,25,27–29]. Shown are Sashimi plots from replicates of "bulk" RNA samples that were isolated from Six2-high and Six2-low mouse fetal kidney cells. Since these cells contain a GFP under the control of a Six2 promoter, and since Six2 is highly expressed in the cap mesenchyme, the Six2-high fraction is predominantly mesenchymal, as it contains mainly cells from the cap mesenchyme (CM), whereas the Six2-low cell fraction contains all other cell populations including all the epithelial cells (PODO, PROX_1, PROX_2, LOH, DIST_CD) as well as some mesenchymal cells from the un-induced mesenchyme (UM). Accordingly, the cassette exons are repressed in the Six2-high cell fraction (that is enriched for the cap mesenchyme, CM) and over-expressed in Six2-low cell fraction (that is enriched with the epithelial cell populations). We isolated total RNA from 3 replicates of cells that were sorted by FACS for high and low levels of the Six2-GFP reporter gene. RNA was isolated using two different kits (TRI reagent and Norgen) and sequenced. We note that the splice isoform switching in Enah and CD44 was difficult to observe in the single cell RNAseq dataset, probably due to the bias and relative sparsity of reads in the single cell data, but was apparent in the "bulk" RNAseq dataset.

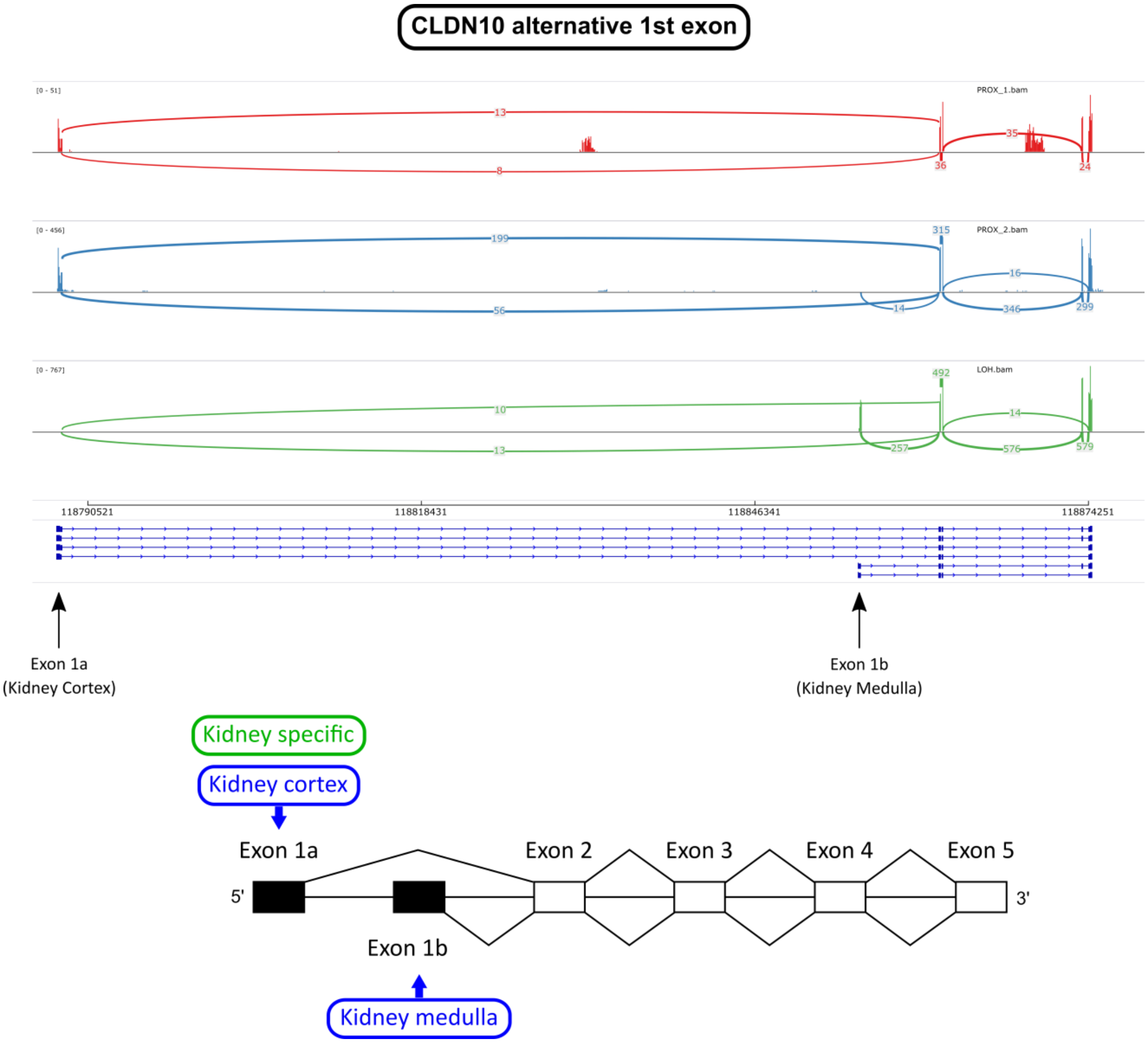


**Figure S19:** Alternative splicing of the gene Cldn10. The isoform Cldn10-1a, a kidney-specific isoform known to be expressed in the kidney cortex [30], is predominantly expressed in the early epithelial structures (PROX_1) and the proximal tubular cells (PROX_2). The isoform Cldn10-1b, which is specific to the kidney medulla, is predominantly expressed in the loop of Henle (LOH). The different isoforms are presumably related to the permeability of the epithelial tight junctions that varies along the different segments of the nephron tubules.

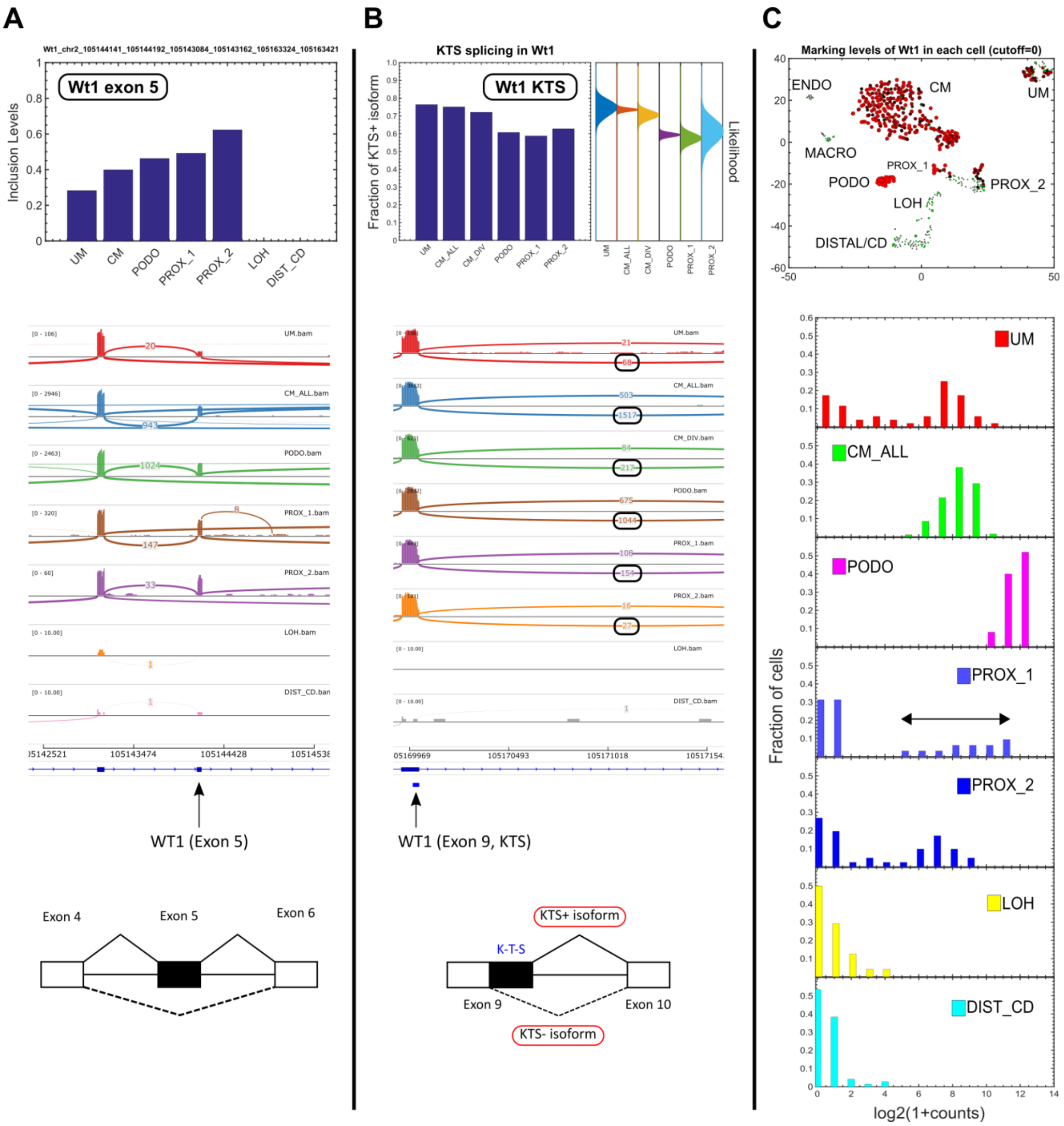


**Figure S20:** Splice isoform switching and gene expression analysis of the gene Wt1, a central gene in kidney development, whose deletion or mutation can lead to developmental defects and pediatric tumors of the kidney (e.g. Wilms' tumors) [31–35]. (A) Cassette exon 5 increases gradually during development. Shown are a barplot of inclusion levels and a Sashimi plot of the *in-silico* "bulk" transcriptomes representing the different populations. (B) KTS alternative splicing in exon 9. Shown are a barplot of inclusion levels, a likelihood plot for the inclusion levels (see Supplementary Information), and a Sashimi plot. An alternative splice donor site at exon 9 inserts or skips the three amino acids lysine (K), threonine (T), and serine (S). The ratio between the Wt1(+KTS) isoform - that includes the KTS segment - and the Wt1(−KTS) isoform – that skips the KTS segment - is typically 60:40 [32,36,37]. A disruption of this ratio to 30:70 is associated with Frasier syndrome, a kidney developmental defect. Here we see that the KTS+:KTS- ratio is approximately 70:30 in the mesenchymal cell populations (the un-induced mesenchyme [UM], the cap mesenchyme [CM], and CM_DIV which consists of only the actively dividing cells within the cap mesenchyme) and converges to approximately 60:40 in the epithelial cell populations (podocytes [PODO], early epithelial structures [PROX_1], and proximal tubules [PROX2]). (C) Single-cell gene expression analysis of Wt1. Shown is a tSNE plot and histograms showing the expression levels of Wt1 in each of the different populations. It can be seen that Wt1 is most highly expressed in the podocytes [38] (PODO), moderately expressed in the un-induced mesenchyme (UM) and cap-mesenchyme (CM), and under-expressed in the loop of Henle (LOH) and distal tubules/collecting duct (DIST/CD). Note the wide distribution of expression in the early epithelial structures (PROX_1), which is probably due to the fact that some cells (e.g. those in the cleft of the S-shaped body [39]) are in the process of differentiating to podocytes while others are destined to become constituents of the proximal tubule, loop of Henle, or distal tubule. The area of each circle in the tSNE plots is proportional to log2(1+expression) of Wt1 in that particular cell.

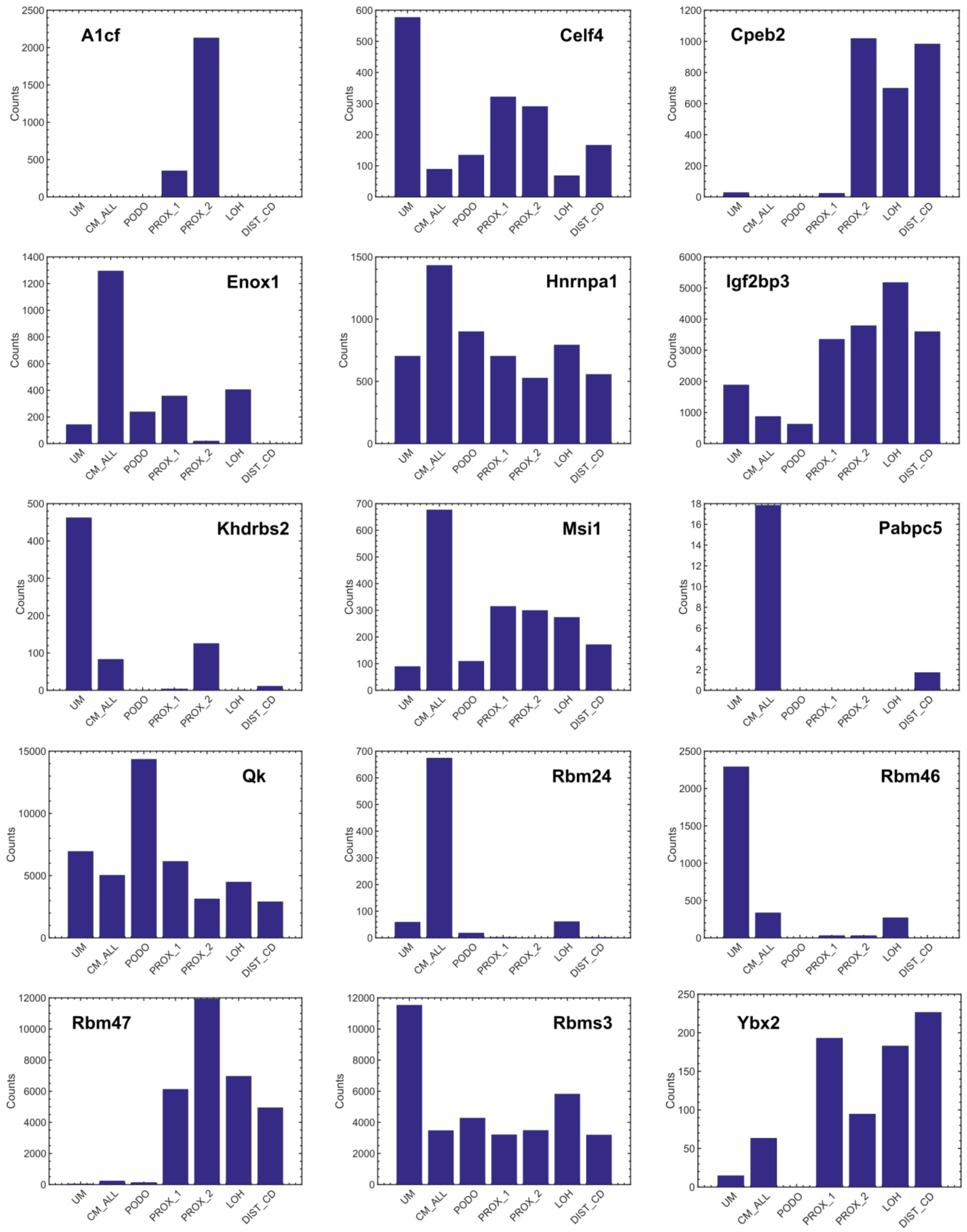


**Figure S21:** Additional putative splicing regulators. Shown are barplots for selected RNA binding proteins (RBP's) [40] that change their expression during the transition from mesenchymal to epithelial states.

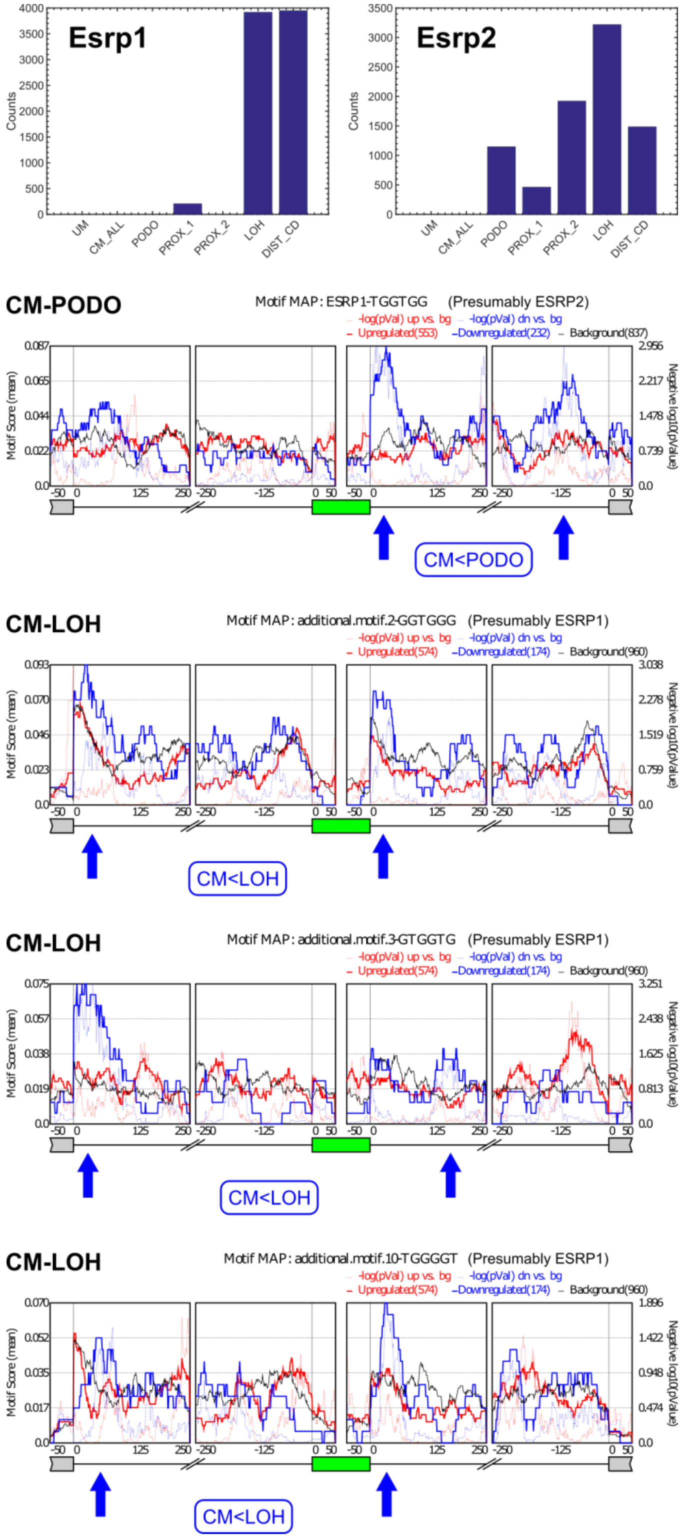


**Figure S22:** Motif enrichment analysis for additional UGG-enriched motifs that were previously found to be binding sites for the RNA binding proteins Esrp1 [26,41] and Esrp2 [42]. Cassette exons that are over-expressed in the epithelial populations contain a significant enrichment of ESRP1/2 binding motifs in their downstream 3’-flanking introns, and in some cases (CM vs. LOH), also in the far 5’ end of their upstream 5’-flanking introns as was previously observed using SELEX-Seq experiments in EMT [41]. This further indicates that ESRP1/2 are splicing regulators involved in Mesenchymal to Epithelial Transition (MET) during kidney development. Note that although the TGGTGG motif (2$^{nd}$ panel from top) is usually regarded as an RNA binding site for Esrp1 [40,41] (Table S4), we hypothesize that in this case it serves as a binding site for Esrp2 since Esrp1 is not expressed in the podocytes and since this motif is similar to the Esrp2 binding motif TGGTG [42,43].
